# Exploring yield stability and the fitness landscape of maize landrace root phenotypes *in silico*

**DOI:** 10.1101/2024.09.07.609951

**Authors:** Ivan Lopez-Valdivia, Harini Rangarajan, Miguel Vallebueno-Estrada, Jonathan P. Lynch

## Abstract

Integrated root phenotypes contribute to environmental adaptation and yield stability. We used the functional-structural plant/soil model *OpenSimRoot_v2* to reconstruct the root phenotypes and environments of eight maize landraces to understand the phenotypic and environmental factors associated with broad adaptation. We found that accessions from low phosphorus regions have root phenotypes with shallow growth angles and greater nodal root numbers, allowing them to adapt to their native environments by improved topsoil foraging. We used machine learning algorithms to detect the most important phenotypes responsible for adaptation to multiple environments. The most important phene states responsible for stability across environments are large cortical cell size and reduced diameter of roots in nodes 5 and 6. When we dissected the components of root diameter, we observed that large cortical cell size improved growth by 28%, 23 % and 114%, while reduced cortical cell file number alone improved shoot growth by 137%, 66% and 216%, under drought, nitrogen and phosphorus stress, respectively. Functional-structural analysis of 96 maize landraces from the Americas, previously phenotyped in mesocosms in the greenhouse, suggested that parsimonious anatomical phenotypes, which reduce the metabolic cost of soil exploration, are the main phenotypes associated with adaptation to multiple environments, while root architectural traits were related to adaptation to specific environments. Our results indicate that integrated phenotypes with root anatomical phenes that reduce the metabolic cost of soil exploration will increase tolerance to stress across multiple environments and therefore improve yield stability, regardless of their root architecture.

## Introduction

The development of crops with greater tolerance to abiotic stress and reduced input requirements is an important goal for global agriculture (Lynch et al., 2023). Improving edaphic resource acquisition by targeted selection of root phenotypes is an opportunity to adapt crops to degraded soil environments (Lynch et al., 2023). The inclusion of root phenotypes into crop breeding programs is challenging due to the complexity of the fitness landscape, phenotyping constraints, and extensive genotype-by-environment interactions (Burridge et al., 2019). However, Burridge et al., (2019) show a successful case where root phenotypes were implemented in breeding programs to significantly improved common bean performance in Mozambique. A better understanding of the fitness landscape of root phenotypes will facilitate the deployment of root phenotypes in breeding programs for more resilient crops.

The functional-structure plant/soil model *OpenSimRoot_v2* (*OSRv2*) is a model that dynamically simulates root growth in three dimensions, soil resource distribution and acquisition, and root-soil interactions (Schäfer et al., 2022). *OSRv2* and its predecessor *SimRoot* allow the parameterization of diverse root phenotypes and environments, permitting analysis of myriad root-soil interactions (Rangarajan et al., 2022). For example, *SimRoot* predicted that root cortical aerenchyma (RCA) would improve plant performance under low fertility soils (Postma and Lynch, 2011), which was then confirmed in field experiments (Saengwilai et al., 2014; Chimungu et al., 2015a; Galindo-Castañeda et al., 2018). Also, *SimRoot* predicted that reduced lateral branching frequency would improve plant performance in low-nitrogen soils but that greater lateral branching frequency would improve plant growth in low-phosphorus soils (Postma et al., 2014; Rangarajan et al., 2018), which was later confirmed in field experiments (Jia et al., 2018; Zhan et al., 2015). It also predicted that the maize/bean/squash polyculture would have greater belowground niche complementarity than the respective monocultures (Postma and Lynch, 2012), which was also confirmed in the field (Zhang et al., 2014). More recently, *OSRv2* capabilities have expanded to include a more robust simulation of photosynthesis, responses to water deficit (Schäfer et al., 2022), and soil impedance (Strock et al., 2022), making it useful in multiple environments.

The ideal integrated phenotype for a particular environment, or “ideotype”, is a function of the circumstances (Rasmusson, 1987). For instance, independent field experiments showed that increased nodal root number (Sun et al., 2018), high lateral branching frequency (Jia et al., 2018), increased root hair formation (Miguel et al., 2015; Zhu et al., 2010a), reduction in root metabolic cost (Galindo-Castañeda et al., 2018; Schneider et al., 2017) and shallow axial root growth angles (Zhu et al., 2005; Miguel et al., 2015) improved the colocalization of roots with phosphorus-rich soil domains, improving crop performance under low phosphorus availability, confirming the utility of ‘topsoil foraging’ ideotype (Lynch and Brown, 2001). In contrast, independent field experiments showed that steep axial root angles (Schneider et al., 2022; Trachsel et al., 2013), fewer nodal roots (Saengwilai et al., 2014a; Gao et al., 2016), few laterals (Zhan et al., 2015; Zhan and Lynch, 2015), and reduction of the metabolic cost of soil exploration (Saengwilai et al., 2014b; Chimungu et al., 2014a; Chimungu et al., 2014b; Jaramillo et al., 2013; Zhu et al., 2010b) colocalized root systems with nitrogen-rich zones in deeper soil regions, improving plant performance in low nitrogen and drought environments, and confirming the utility of the ideotype ‘steep, cheap and deep’ (Lynch, 2013).

It has been proposed that genotypes that react less to the environment are needed to guarantee consistent crop production in a range of environments (Francis and Kannenberg, 1978). Most yield stability studies have tested several hybrids in multiple environments and normally particular hybrids are presented as the most stable (e.g., Shiri et al 2013; Zhao et al., 2018; Fan et al., 2007), however, the factors underlying yield stability are not well understood. It has been proposed that yield stability is associated with increased stress tolerance (Tollenaar and Lee, 2002). On the other hand, it has been demonstrated that soil environments that reduce stress, for instance, by increasing soil water holding capacity, are associated with yield stability (Williams et al., 2016). More recent studies have shown that optimal fertilization (Ahrends et al., 2020; Knapp et al., 2018) often results in greater yield stability. Previously, we described how specific combinations of root anatomical and architectural phenotypes improve soil exploration in specific edaphic environments, improving plant performance (Lynch et al., 2023; Lynch 2022). However, it is unknown if integrated root phenotypes, that increase tolerance to edaphic stress, would also contribute to yield stability.

Root architectural phenotypes that colocalize roots with soil resource availability improve plant performance in soils with suboptimal water, nitrogen, and phosphorus availability (Lynch et al., 2023), while root anatomical phenotypes that reduce the metabolic cost of soil exploration will improve plant performance under nitrogen, water, and phosphorus stress (Lynch 2022). Therefore, we hypothesize that anatomical phenotypes that reduce the metabolic cost of soil exploration and intermediate architectural phenotypes will be broadly adapted to multiple environments. In this study we used *shovelomics* and Laser Ablation Tomography (LAT) to characterize the root anatomy and architecture of 8 maize landraces in the field. We used *OSRv2* to reconstruct the root phenotypes and their native soil and atmospheric environments *in silico*. We evaluated their performance and used machine learning models to determine the most relevant phene states associated with environment-specific and broad adaptation. We constructed synthetic root phenotypes designed for environment-specific and overall optimal plant performance. Furthermore, we extended our analysis to 96 maize landraces, previously phenotyped by Burton et al., 2013, by simulating their growth in the 8 environments using OSRv2 and evaluating the most important phenotypes for broad adaptation. Finally, we evaluated the components of the synthetic phenotypes independently to determine their contributions to nitrogen and phosphorus stress.

## Material and Methods

### Model description

*OSRv2* is a functional-structural plant/soil model that simulates root growth in three dimensions, soil nutrient distribution, and root-soil interactions (Postma et al., 2017; Schäfer et al., 2022). Root architecture is simulated as a hierarchical development of root classes, branching frequencies, growth vectors, and growth rates. *OSRv2* includes a carbon module that calculates the available carbon for growth based on a subtraction of the carbon sources minus the carbon sinks. The carbon sinks are the root exudates and the maintenance cost due to respiration, while the carbon sources are the carbon available in the seed and the carbon produced by photosynthesis. Carbon partitioning rules determine that exudates and respiration are obligate costs, once this demand is covered the remaining carbon could be used for growth. The model considers constraints to root and shoot growth due to the availability of resources such as nitrogen, phosphorus, and water (Schäfer et al., 2022), but also due to soil impedance (Strock et al., 2022). The shoot is simulated by several state variables in a non-geometrical fashion (Schäfer et al., 2022). Responses to water deficit are simulated considering Farquhar-von Caemmerer-Berry model for photosynthesis, leaf gas exchange and stomatal conductance, leaf temperature and energy balance models, sun/shade model for leaf-to-canopy scaling, a nitrogen-limited photosynthesis model, water stress response functions and models for day/night cycles (Schäfer et al., 2022) making it sensitive to fluctuations in atmospheric conditions such as CO_2_, temperature, solar radiation among others. The movement of water in the soil is simulated by the SWMS3D model (Simunek, 1995). The radial and axial conductivity of water in the roots is simulated by a hydraulic network model (Alm et al., 1992; Doussan et al., 1998). Evapotranspiration is simulated with the Penman-Monteith equations (Penman 1948; Monteith 1965). Soil hydraulic properties, root water uptake, precipitation, and evaporation will determine the soil water dynamics (Schäfer et al., 2022). Nitrogen movement in the soil is also simulated by the SWMS3D model. Nitrogen mineralization is simulated by the Yang and Janssen model (Yang & Janssen, 2000). To simulate plant response to stress the model calculates stress factors based on the optimal resource concentration in the tissue versus the actual resource acquisition. The stress factors will directly impact leaf expansion rate and light use efficiency. When the leaf expansion rate is constrained, the sink of the shoot is reduced, and proportionately more carbon is allocated to the roots, functioning as a growth regulator. On the other hand, when light use efficiency is affected, there is less carbohydrate availability and a reduction of growth in general. The impact of stress factors on leaf expansion rate and light use efficiency is resource-specific (Postma, 2011; Schäfer et al., 2022).

### Accession selection

We selected 8 maize (*Zea mays L. subsp. mays*) accessions from a root phenotypic diversity panel (Burton et al., 2013) from the most extreme environments in terms of aridity and humidity (Table S1); 4 accessions from humid regions (Costa_Rica [PI471823], Cusco, Peru [PI571629], Huanuco, Peru [PI571913], Madre de Dios, Peru [PI571994]), and 4 accessions from arid regions (Mendoza, Argentina [PI516002], Lambayeque, Peru [PI571439], Ica Peru [PI571523], Colorado, United States [PI608619]). Arid regions also coincided with soils having low nitrogen availability but high in phosphorus availability, while humid regions tend to have soils with greater nitrogen availability but low phosphorus availability. The location of every accession was obtained from georeferenced information available at USDA, Agricultural Research Service, National Plant Germplasm System (2024). We obtained soil nitrogen, phosphorus, and precipitation data from SoilGrids (Poggio et al., 2021), a dissertation by Jaramillo-Velasteguí (2011), and POWER Data Access Viewer v2.0.0 (https://power.larc.nasa.gov/data-access-viewer/ consulted on 2023/09/15), respectively.

### Root parameterization

#### Architecture

We grew 10 plants per accession at the Russell E. Larson Agricultural Research Farm at Rock Spring, PA, USA (40°42’40.915” N, 77°,57’11.120” W) from June through September 2021. We fertilized using 150 kg N ha^-1^ (Urea) to fulfill the optimal requirements for maize considering soil tests before planting. We provided drip irrigation as needed. Planting was performed in single-row plots at a density of 73,300 plants ha^-1^. After 90 days of growth, all plants were sampled following the shovelomics procedure (Trachsel et al., 2011). Roots were washed and processed for subsequent phenotyping. Root phenotyping consisted of imaging the root system while sequentially removing a set of nodal roots per node. This process was repeated in every node producing an image for every node until we removed all the nodal roots. Once we removed all nodal roots, we took a final image of the seminal roots and counted the seminal root number manually. We used ImageJ (Sosa et al., 2014) to analyze the images. For every node, we measured the angle, nodal root number, lateral branching frequency, and root diameter. The root angle was measured considering the horizon as the starting point. For every accession, we integrated their root phenotypic data per node into *OSRv2* using the input file production script available in the following link (https://github.com/ilovaldivia/LopezValdivia_2024_YieldStability.git).

#### Anatomy

To measure root anatomy we sampled 4 cm root segments, 5 to 9 cm from the base of a second-node root for all plants and we stored them in 70% (v/v) ethanol in water. We imaged three cross-sections along the 4 cm root segment utilizing Laser Ablation Tomography (LAT) (Strock et al., 2019). LAT consists of an oscillating pulsed laser (s-Pulse HP, 343 nm THG, Amplitude Systems) that segments the root sample while a camera (α7R III digital camera, Sony. MP-65 mm F/2.8 Lens photo macro 1–5, Canon) gathers cross-sectional images of the ablated root surface. We measured cortical cell size (CCS), cortical cell file number (CCFN), and root cortical aerenchyma (RCA) using RootScan 2.4v (https://plantscience.psu.edu/research/labs/roots/methods/computer/rootscan). We used a multiple linear regression model reported by Lopez-Valdivia et al., (2023) to predict respiration and nitrogen construction cost per gram of root tissue using CCS and CCFN as predictors. We integrated CCS and CCFN into *OSRv2* as root respiration and nitrogen construction cost predictions, while RCA was integrated directly.

### Environment Parameterization

We designed 8 environments in *OSRv2* corresponding to the origin of each accession. To design the environments, we obtained soil and atmospheric parameters from SoilGrids (Poggio et al., 2021) and POWER Data Access Viewer v2.0.0 (https://power.larc.nasa.gov/data-access-viewer/ consulted on 2023/09/15), respectively. For the atmosphere, we extracted data for wind speed, minimum and maximum temperature, precipitation, and relative humidity in a period from 1980 to 2021 (Data S1). For each atmospheric parameter, we used an average of the 1980-2021 period, and we selected 60 days corresponding to a growing season considering the maize planting dates in each region (Data S2). For the soil, we used soil core information from SoilGrids to extract nitrogen concentration, soil texture, bulk density, and soil organic matter. The closest core to the coordinate of origin was considered a representative of the soil environment. Soil core IDs and soil core coordinates are available in Data S3. We used the soil texture to calculate the van Genuchten parameters (Data S4) utilizing Rosetta3 (https://www.handbook60.org/rosetta/). We inferred the soil taxon from the USDA taxonomy system using soil texture information and FAO classification, and we obtained the average phosphorus concentration for that soil class from Jaramillo-Velasteguí (2011). We integrated the soil and atmospheric data into *OSRv2* with the input file production script available at: https://github.com/ilovaldivia/LopezValdivia_2024_YieldStability.git.

### Model runs

To test the performance of the 8 accessions across the 8 environments we ran 64 simulations of all accession/environment combinations in *OSRv2* (Schäfer et al., 2022) with git version 9621150b6f. To test the performance of synthetic phenotypes we ran 72 simulations for 9 synthetic phenotypes (8 environment-specific and 1 general) under 8 environments. Furthermore, we ran 12 simulations for the dissected phenotypes which included the independent effect of branching frequency (BF), cortical cell size (CCS), and root diameter of nodes 5 and 6 (Dia), as well as combinations of them (BF+CCS; BF+Dia; CCS+Dia). A reference root phenotype with contrary values of BF, root diameter, and CCS compared with the general synthetic was simulated for comparison proposes. Simulations were run utilizing High-Performance Computing at Pennsylvania State University. All input files required to replicate the simulations can be found here: https://github.com/ilovaldivia/LopezValdivia_2024_YieldStability.git. All data was extracted and analyzed using R version 4.2.3 (R Core Team, 2019).

### Synthetic phenotypes

To identify the most important root phenotypes in each environment and across all environments we ran a Boruta feature selection algorithm in R version 4.2.3 (R Core Team, 2019). We used the most important root phenotypic parameters for each environment to build a multiple regression model using R version 4.2.3 (R Core Team, 2019). We used the coefficients of the multiple regression to modify the variables that were selected in Boruta. For instance, if the coefficient was positive for the branching frequency for node 4, we would increase the value of the synthetic phenotype to the maximum value available in the distribution of the 8 maize accessions. We repeated the process for all relevant phenotypes in all environments to create the specific synthetic phenotypes. To create the general synthetic root phenotype we repeated the process using the average of all the environments.

### Functional analysis of maize populations of the Americas

To explore the functional utility of root phenotypes across the Americas we analyzed root anatomy and architecture data available from Burton et al., (2013) using a combination of machine learning algorithms and the functional-structural model OpenSimRoot_v2. We selected the data only for landraces of maize which had geo-localization data. We performed a principal component analysis and k-means clustering to find the main clusters of phenotypes using R Studio version 4.2.3 (R Core Team, 2019) and the MASS package (Venables and Ripley, 2002). To evaluate a correlation between geographic location and the root phenotypic clusters, we compared the latitude variances among the clusters using one-way ANOVA and Tukey significance test. We classified the main clusters as categorical variables and we built a linear discriminant model to predict the cluster category based on environmental data including precipitation, elevation, nitrogen soil concentration and phosphorus soil concentration. Finally, we used *OpenSimRoot_v2* to simulate root growth corresponding to the clusters of root phenotypes identified by the principal component analysis in the 8 environments described earlier in the ‘environment parameterization’ section. Simulation data was analyzed and graphed using R Studio version 4.2.3 (R Core Team, 2019) and the ggplot2 package (Wickham, 2016).

## Results

### Variation of root phenotypes across multiple nodes

Root phenotypes display wide variation among open-pollinated populations (Burton et al., 2013). We observed no variation in the number of nodal roots in nodes 1, 2, and 3, while significant variation was observed for nodes 4, 5, and 6 (Figure 1). Root anatomy showed less variation for cortical cell file number (8-12), and percentage of aerenchyma (5-25%) compared with the phenotypic ranges observed by Burton et al., (2013) in 195 landraces from the Americas (6-15 and 6-37% for cortical cell size and percentage of aerenchyma, respectively). However, we observed different phenotypic variation when compared with data from inbred populations. For instance, we observed that cortical cell size in landraces (393-1434 µm) was generally larger than in inbred IBM populations (100-900 µm^2^) as reported in Chimungu et al., (2014) and Lopez-Valdivia et al., (2023). Overall, our root phenotypic measurements represent sufficient phenotypic variation for the purposes of this study.

**Figure 1.**
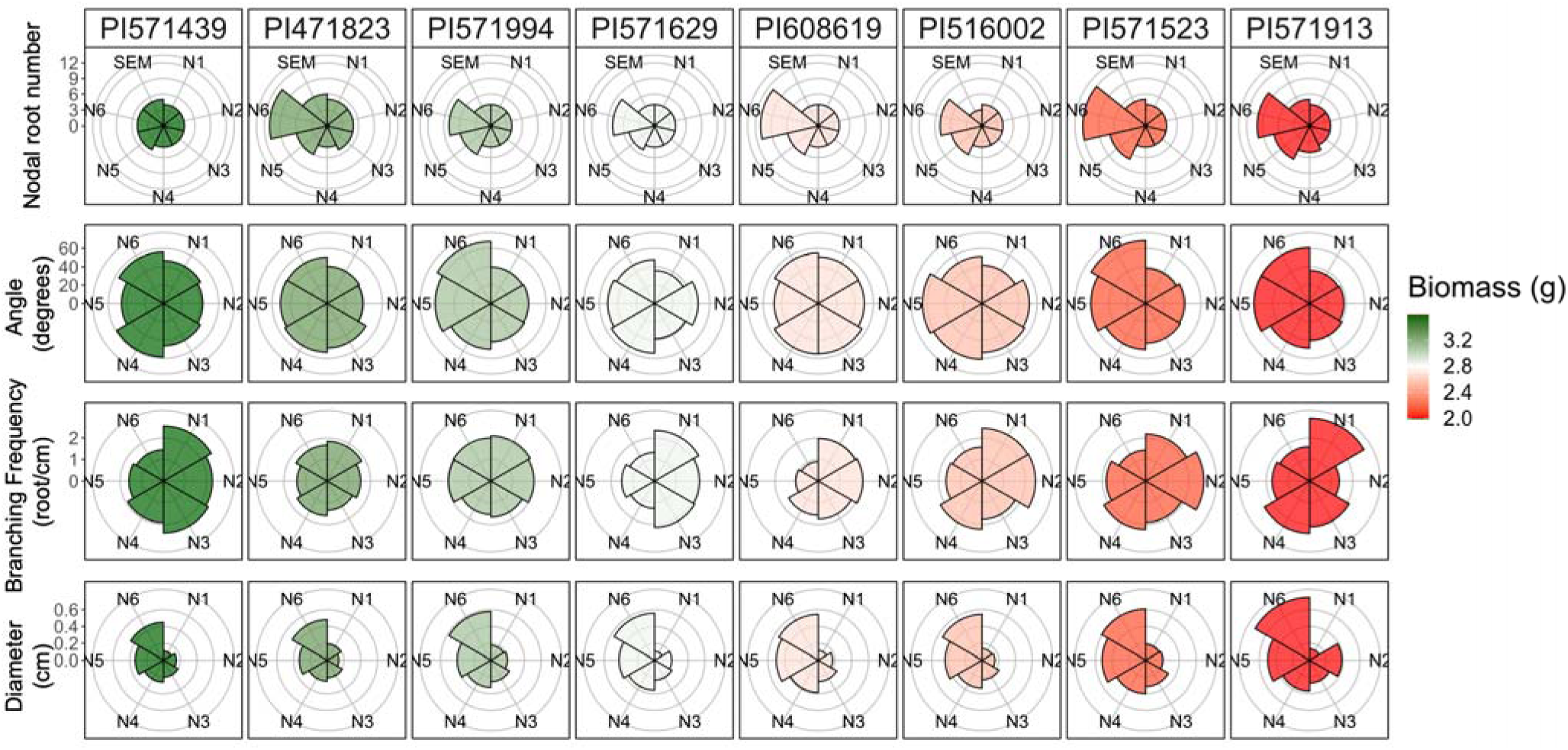
Parameterization of the root system architecture of 8 maize landrace accessions coming from the Americas. Accessions are organized from left to right from the greatest to the lowest average shoot biomass across the 8 environments. All measurements were taken by node and ‘N’ represent the node number. SEM represents seminal roots.

### Non-stable accessions showed contrasting strategies for soil exploration

The spatial and temporal colocalization of roots and soil resources will determine resource uptake and its utility in a particular environment. To explore the utility of each of the 8 integrated phenotypes we reconstructed their root phenotypes and environments in *OpenSimRoot*, and we simulated a reciprocal transplanting (we will refer to the native environments as the accession name in italics plus the letter ‘e’ at the end, E. g. *PI571629e*). We observed that the rankings of 6 accessions were stable across all environments. In contrast, 2 accessions (PI471823 and PI571994) ranked between the first or second best in some environments but then shifted to fourth to sixth place in other environments. We classified the consistent and non-consistent ranking accessions as stable and non-stable, respectively (Figure 2). To analyze the differences between the non-stable accessions we compared their root phenotypes and evaluated their performance in contrasting environments. Accession PI471823 has shallower root angles in younger nodes, greater number of nodal roots, and reduced lateral root branching frequency compared with PI571994 (Figure 3), including some of the phenotypes of the ‘topsoil foraging’ ideotype (Lynch and Brown, 2001). Accession PI471823 also had greater shoot biomass compared with accession PI571994 in environments *PI571629e* and *PI471823e*, which showed the greatest levels of phosphorus stress. On the other hand, accession PI571994 had steeper angles, a reduced number of nodal roots, and increased lateral branching frequency (Figure 3). Steeper root growth angles and reduced production of nodal roots are elements of the ‘steep, cheap and deep’ ideotype (Lynch, 2013). The simulation of the root phenotype of PI571994 also produced greater biomass in environments *PI571439e*, *PI516002e*, *PI608619e*, *PI571629e*, which have the greatest nitrogen stress. Overall, the non-stability of accessions PI471823 and PI571994 is due to their extreme root phenotypes, especially regarding nodal root number and root growth angle, improving their adaptation to specific environments but making them unstable across all the environments.

**Figure 2:**
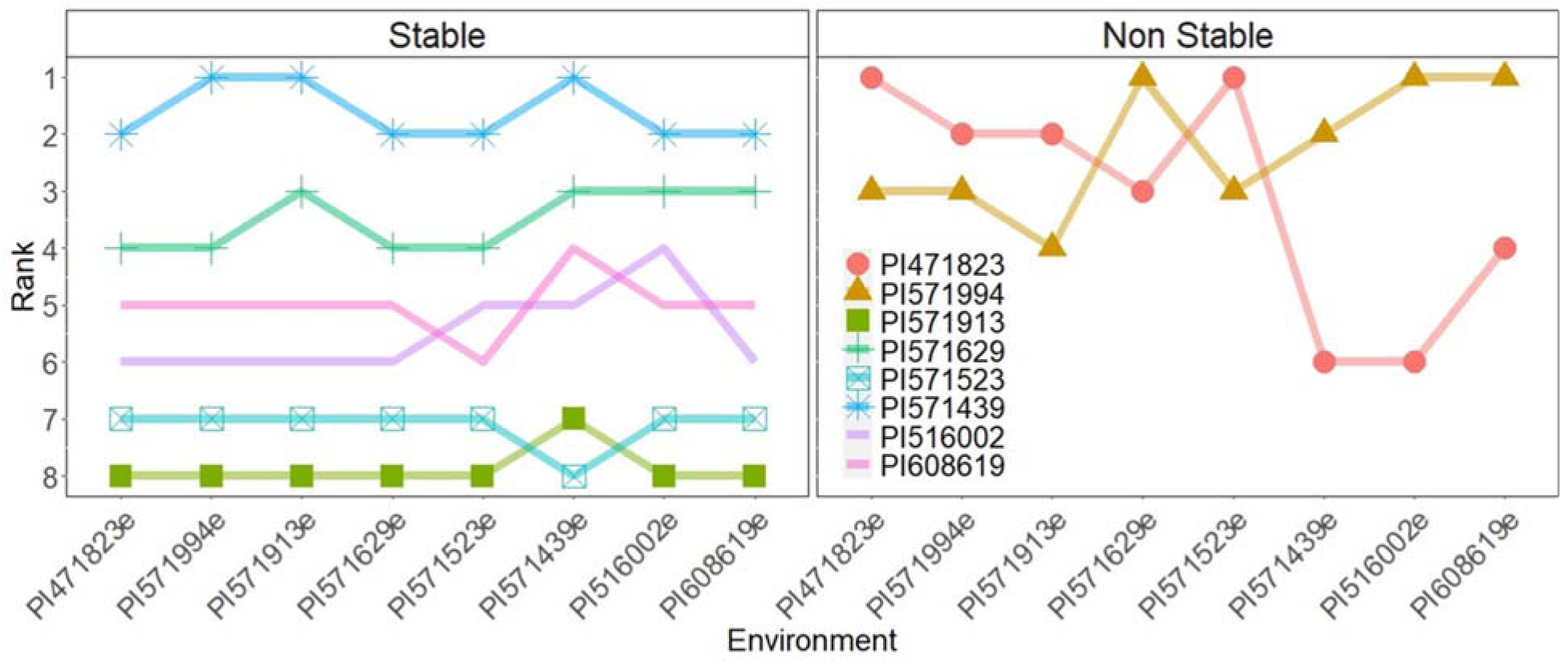
Simulated reciprocal transplanting of 8 maize landraces under 8 environments. Accessions are divided into 2 groups corresponding to their pattern across all environments. Stable (PI571629, PI571439, PI571913, PI516002, PI608619) and non-stable (PI471823, PI571994).

**Figure 3:**
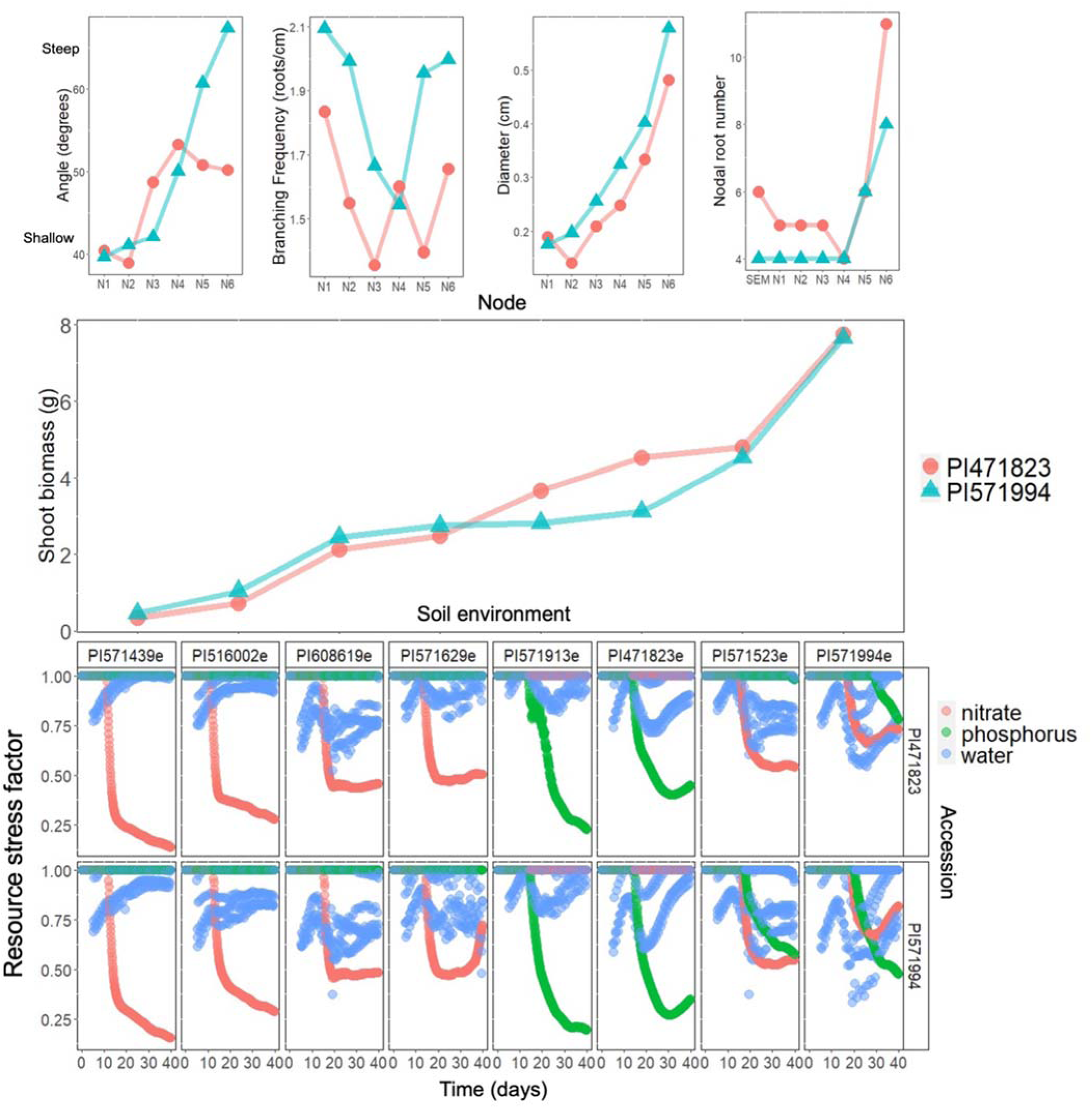
Non-stable maize landraces have contrasting integrated root phenotypes. PI571994 shows steeper angles, less branching frequencies, thicker root diameters, and fewer nodal roots compared with PI471823. Top panels represent the root phenotypes per node and ‘N’ represent the node number. Middle panel presents shoot biomass produced in the 8 environments. Bottom panel represents the nitrate, phosphorus, and water stress factors in the soil in a 40-day simulation, where stress factor values closer to 0 represent greater stress.

### Phene states associated with greater stability include large cortical cell size

To understand root phenotypes contributing to stability across environments we employed a Boruta feature selection algorithm to find the most important phenotypes for each environment and across all environments. In general, lateral branching frequency in node 4, cortical cell size, and root diameter of nodes 5 and 6 were important across all environments (Table 1). Every environment had a specific combination of phenotypes that were important. For instance, in environments *PI471823e* and *PI571913e,* root diameters are the most important variables determining biomass (Table 1), while in environments *PI516002e* and *PI571439e*, nodal root number and cortical cell size are the most important variables determining biomass (Table 1). To understand if an optimized root phenotype for a specific environment would surpass a root phenotype optimized for multiple environments, we designed synthetic phenotypes for specific and general environments using multiple linear regression on the root phenotypes selected previously by Boruta (a detailed description of the synthetic phenotypes design is available in the methods section). We observed that the general synthetic root phenotype produced greater biomass compared with the original phenotypes in 6 of 8 environments (Figure 4). Most of the specific synthetic phenotypes were superior to their original (Figure 4), except for accession PI471823. However, the general synthetic root phenotype produced greater biomass than 5 of the specific synthetic phenotypes. The synthetic phenotypes customized for environments *PI516002e*, *PI608619e,* and *PI571439e* had similar performance as the general synthetic phenotype (Figure 4). The general synthetic phenotypes and synthetic phenotypes customized for environments *PI516002e*, *PI608619e,* and *PI571439e* share a large cortical cell size, which is likely the factor driving their superior performance. The general synthetic phenotype had the best performance across environments (Figure 4). Synthetic phenotypes generated for their specific environments outperformed the actual phenotype from that environment in almost all cases (Figure 4).

**Figure 4:**
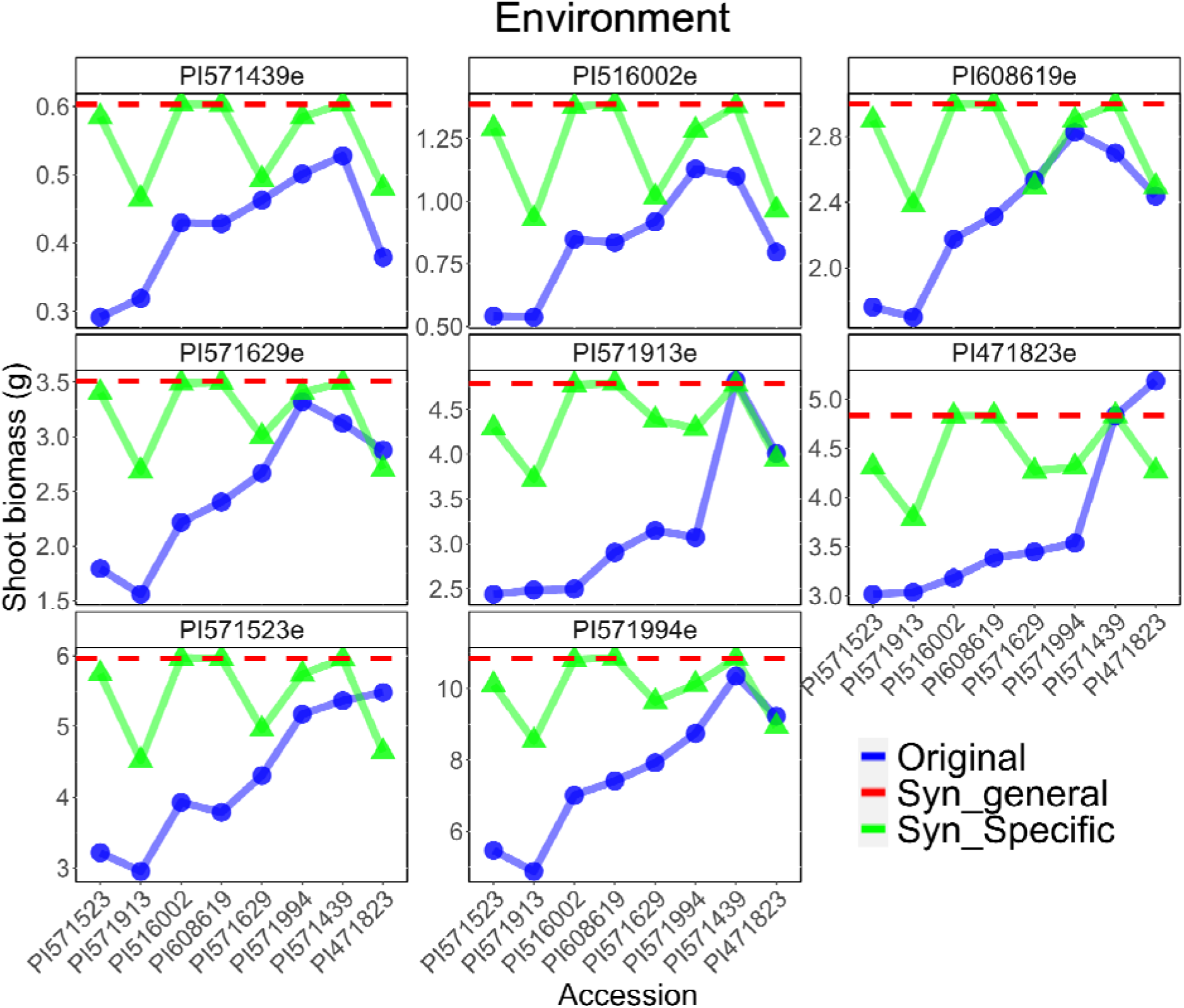
Shoot biomass of synthetic root phenotypes designed for specific and general environments. Environments are organized from top-left to bottom-right from the lowest to the greatest shoot biomass production. The blue line (Original) represents the original root phenotype for each accession. The green line (Syn_Specific) represents the specific synthetic phenotype that was optimized for its native environment. The red line (Syn_general) represent the general synthetic phenotype that was optimized for all environments. Synthetic phenotype are described in Table 2.

**Table 1.**
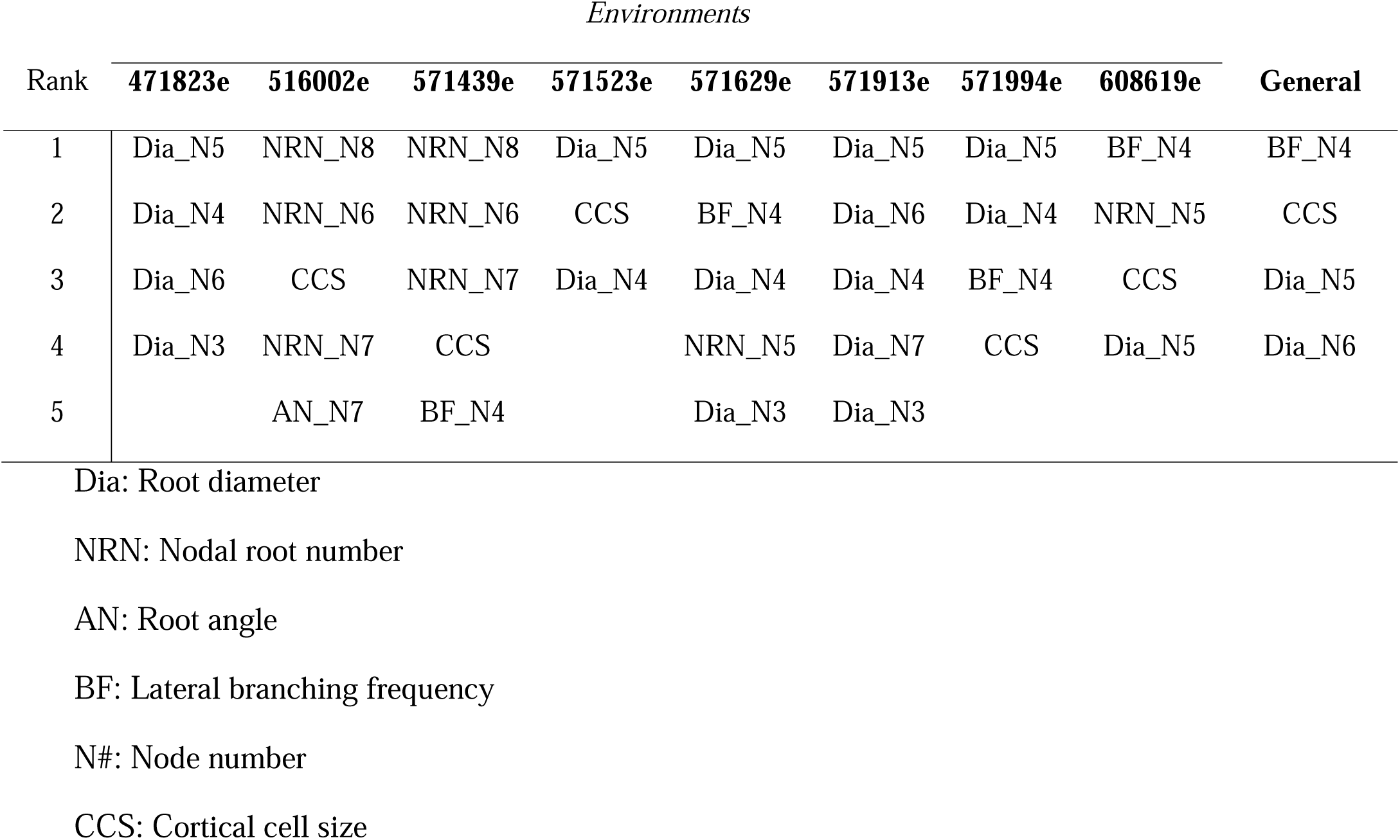
Feature selection with Boruta of the root phenotypes that are important in every environment and across all environments (General).

| *Environments* |  |  |  |  |  |  |  |  |  |
| --- | --- | --- | --- | --- | --- | --- | --- | --- | --- |
| Rank | 471823e | 516002e | 571439e | 571523e | 571629e | 571913e | 571994e | 608619e | General |
| 1 | Dia_N5 | NRN_N8 | NRN_N8 | Dia_N5 | Dia_N5 | Dia_N5 | Dia_N5 | BF_N4 | BF_N4 |
| 2 | Dia_N4 | NRN_N6 | NRN_N6 | CCS | BF_N4 | Dia_N6 | Dia_N4 | NRN_N5 | CCS |
| 3 | Dia_N6 | CCS | NRN_N7 | Dia_N4 | Dia_N4 | Dia_N4 | BF_N4 | CCS | Dia_N5 |
| 4 | Dia_N3 | NRN_N7 | CCS |  | NRN_N5 | Dia_N7 | CCS | Dia_N5 | Dia_N6 |
| 5 |  | AN_N7 | BF_N4 |  | Dia_N3 | Dia_N3 |  |  |  |

Dia: Root diameter
NRN: Nodal root number
AN: Root angle
BF: Lateral branching frequency
N#: Node number
CCS: Cortical cell size

**Table 2:**
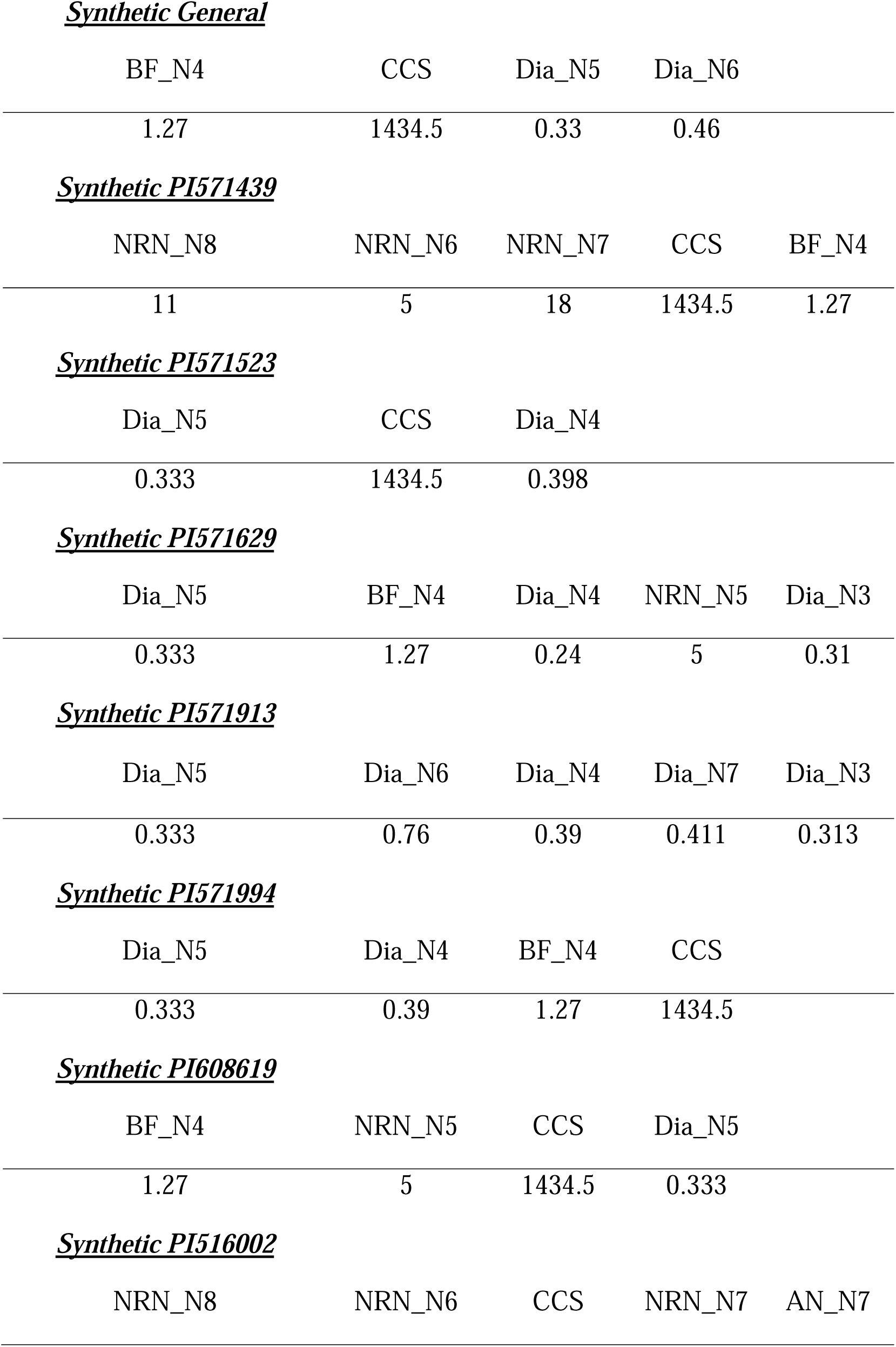
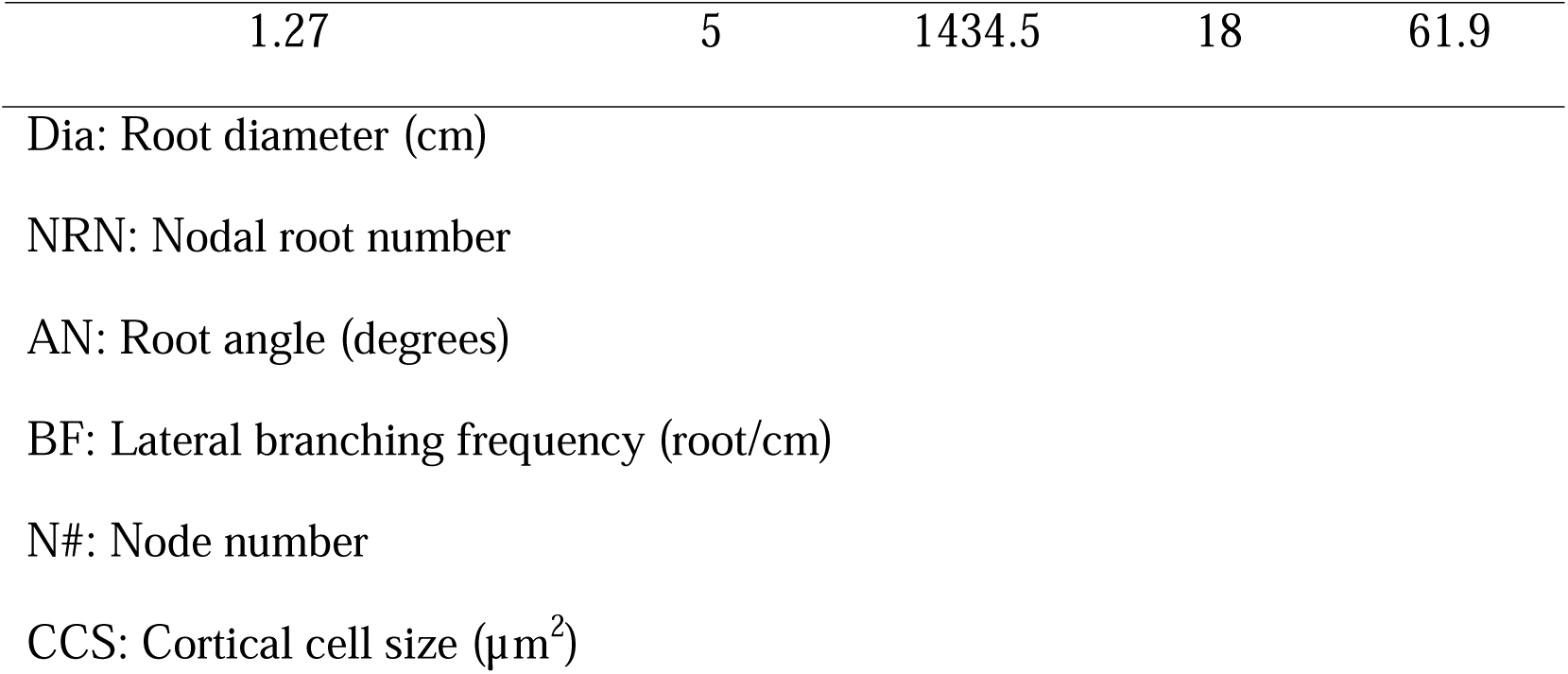
Synthetic integrated phenotypes built based on Boruta and multiple linear regression.

### Evaluation of individual components of synthetic phenotypes

To understand the individual contributions of each phene state to the general synthetic phenotype we simulated the effect of each phene state alone and evaluated its performance compared with a reference phenotype. The effect of reduced root diameter in nodes 5 and 6 (Dia) improved shoot growth 3, 13 and 19% compared with the reference, in drought, nitrogen and phosphorus stress conditions, respectively (Figure 5). Large CCS improved biomass 28%, 23 % and 114% compared with the reference phenotype in drought, nitrogen and phosphorus stress, respectively. The combined effect of large CCS and smaller diameter (synthetic) showed an additive effect by improving shoot growth 35%, 33% and 134% compared with the reference under drought, nitrogen and phosphorus stress, respectively (Figure 5). We observed a minor increase in biomass of decreased branching frequency of axial roots of node 4 of 0.54 and 0.4% compared with the reference in nitrogen and phosphorus stress, respectively. Decreased branching frequency of axial roots of node 4 decreased plant biomass by 1.86% under drought. Overall, the greatest effect of the synthetic phenotypes is given by large CCS and reduced diameter of nodes 5 and 6, while the effect of branching frequency in node 4 is irrelevant.

**Figure 5:**
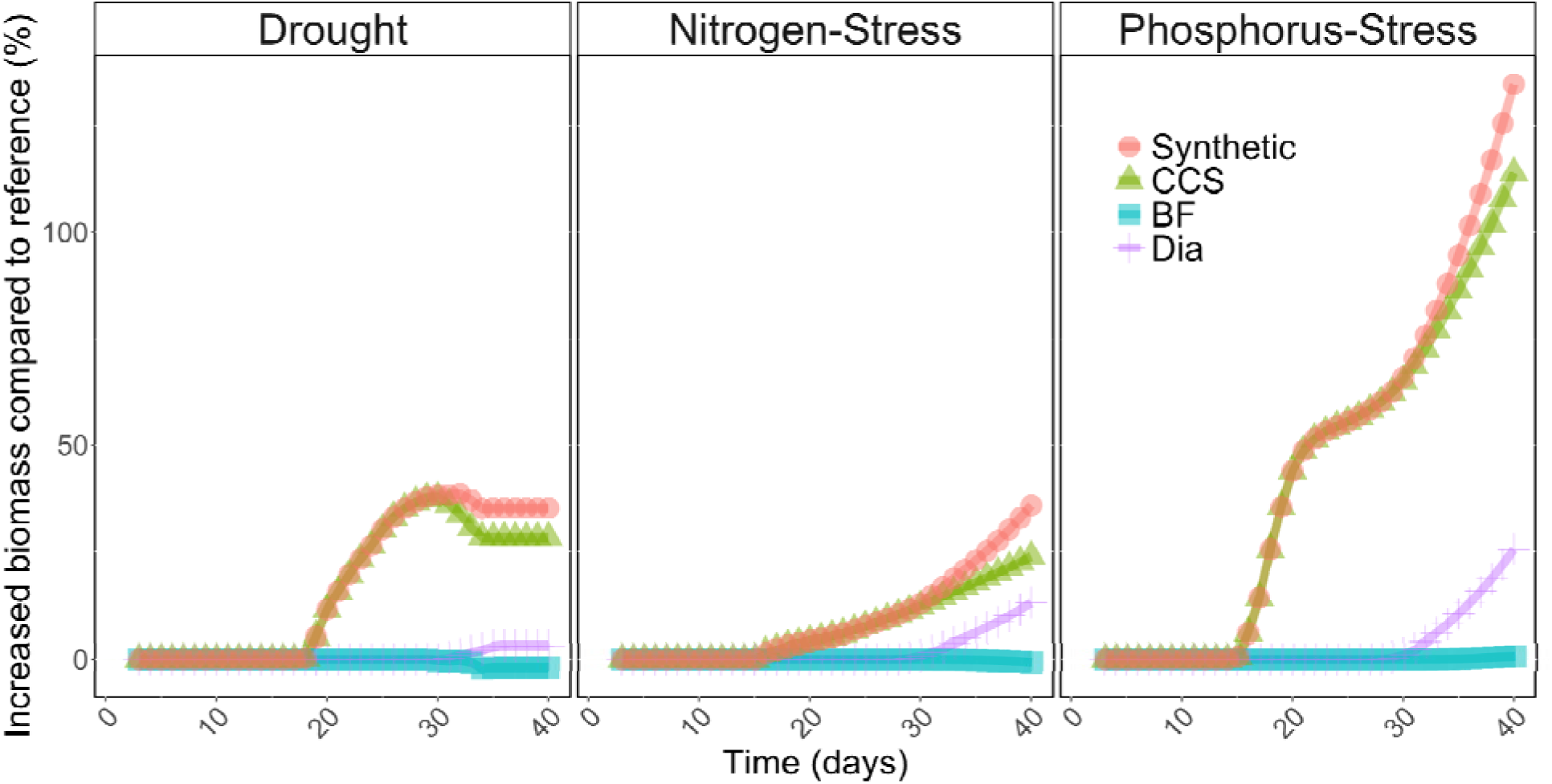
Evaluation of each phene state of the Synthetic phenotype under drought in isolation (left panel), nitrogen stress in isolation (center), phosphorus stress in isolation (right panel). The ‘Synthetic’ phenotype represents all phene states simulated together. CCS represents the independent effect of large cortical cell size, BF represents the independent effect of reduced lateral branching frequency, Dia represents the independent effect of small root diameter of nodes 5 and 6.

### Functional characterization of root phenotypes of maize landraces

#### Principal Component analysis and K-means clustering of root phenotypes

There is extensive variation for root phenotypes among landraces of maize. To capture greater phenotypic diversity in our analysis we performed a principal component analysis and K-means clustering in a maize diversity panel (Burton et al., 2013) to find clusters of root phenotypes. We found 4 main clusters (Figure 6A) distributed across different regions of the Americas (Figure 6B). Clusters 1 and 2 showed significative differences in their spatial distribution in the Americas (Table 3; Figure 6B). Cluster 1 distributed mostly in North America, while cluster 2 mostly distributed in the Andean highlands of South America (Figure 6B, Figure 7). The root phenotypes grouped in cluster 1 were characterized by having shallow nodal growth root angle, thick nodal roots, with large anatomical phenotypes (E.g. large xylem vessels, large stele area, increased cortical cell file number, and cortical cell area; Figure 7). In contrast, root phenotypes grouped in cluster 2 tend to have steeper root growth angles, fewer nodal root number and thin nodal roots with smaller anatomical phenotypes (e.g. small xylem vessel area, reduced cortical cell file number, small stele area and reduced cortical cell area; Figure 7). Clusters 3 and 4 distributed evenly across the Americas. The root phenotypes grouped in cluster 3 were characterized by having an average phenotype (Figure 7), while root phenotypes classified in cluster 4 tend to have a greater number of thin nodal roots with shallow growth angle and small anatomical phenotypes (E. g. small xylem vessel area, small stele area, reduced cortical cell files number and small cortical cell area; Figure 7).

**Figure 6:**
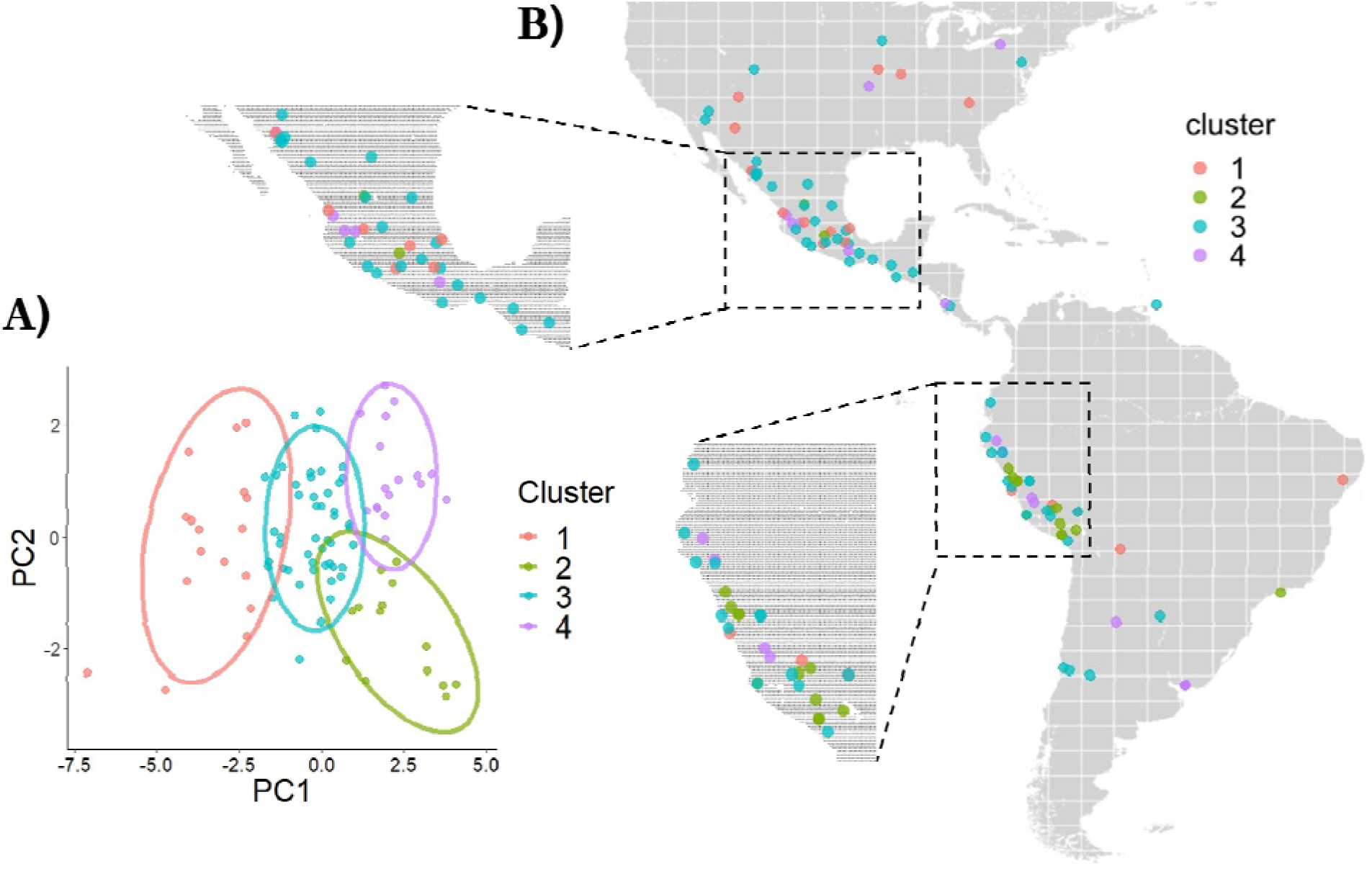
Principal component analysis and K-means clustering of root phenotypes from 100 maize landraces of Burton (2013) panel. A) Shows k-means clustering in 4 main groups. B) Geographic distribution of maize landraces across the Americas.

**Figure 7:**
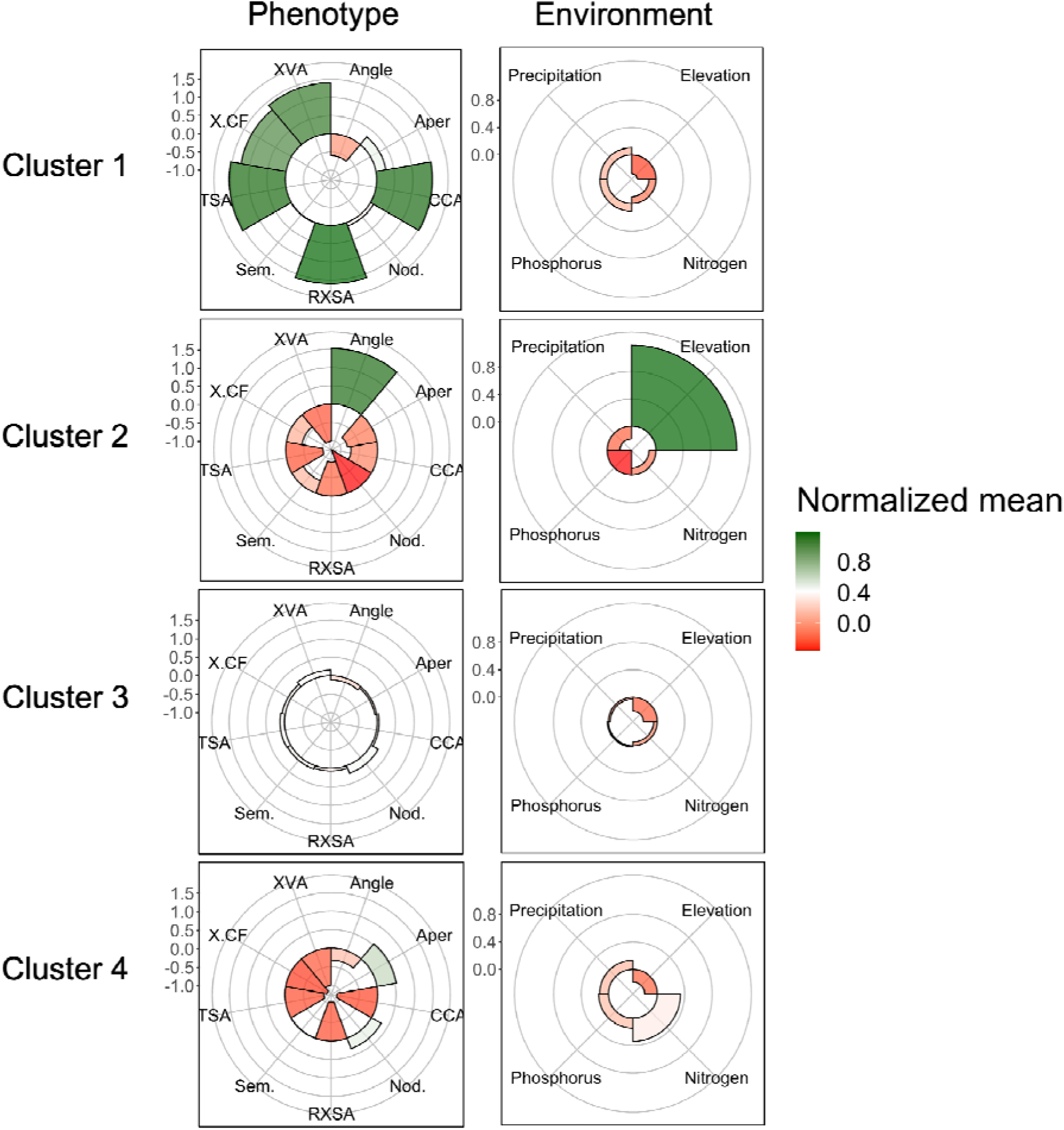
Normalized values of root phenotypes and environmental factors describing the 4 clusters identified by PCA and K means. Root phenotype abbreviations correspond to XVA, xylem vessel area; X.CF cortical cell file number; TSA, total stele area; Sem. Seminal root number; RXSA, root cross-section area; Nod. Nodal root number; CCA, cortical cell area; and Aper, percentage of aerenchyma.

**Table 3.** Analysis of variance and Tukey test of the importance of latitude among 4 clusters identified by PCA and k-means. Comparison of cluster means are shown in the cluster column. For instance, 2-1 refers to the comparison of cluster 1 and cluster 2 means.

| Cluster | Difference | lwrLower | Upper | p adj |
| --- | --- | --- | --- | --- |
| 2-1 | -24.2663 | -44.3459 | -4.18667 | 0.011212 |
| 3-1 | -7.12959 | -22.8323 | 8.573106 | 0.635918 |
| 4-1 | -15.5939 | -34.6772 | 3.489367 | 0.148837 |
| 3-2 | 17.13668 | 0.23718 | 34.03618 | 0.045577 |
| 4-2 | 8.672351 | -11.4072 | 28.75195 | 0.671965 |
| 4-3 | -8.46433 | -24.167 | 7.238367 | 0.496121 |

#### Linear discriminant analysis captures environmental variables

Cluster 1 and 2 were distributed in North and South America, while cluster 3 and 4 distributed evenly across the Americas. To understand the environmental variables associated with each cluster we performed a linear discriminant analysis using the cluster 1-4 as categories and environmental variables as predictors. Cluster 1 was described mainly by lowlands in North America with less availability of nitrogen, but greater availability of phosphorus and precipitation compared with other clusters (Figure 7). Cluster 2 was described by the highlands of South America, with low availability of phosphorus and nitrogen, and reduced precipitation (Figure 7). Cluster 3 was described by lowlands and average conditions for phosphorus, nitrogen and precipitation (Figure 7). Cluster 4 was described by lowlands and relatively abundant nitrogen conditions. Our model predicted the cluster category in the test dataset with 58% accuracy. Phenotypes grouped in cluster 1, with shallow and thick nodal roots, come from low nitrogen environments, while phenotypes grouped in cluster 2, with fewer and steep nodal roots, tend to have reduced precipitation and low phosphorus soils.

#### Functional analysis of root phenotypes using *OSR_v2*

To understand the functional utility of the integrated phenotypes grouped in cluster 1-4, we simulated the root growth of each cluster in 8 contrasting environments using *OSR_v2*. We observed that the root phenotype representing cluster 4 produced a greater biomass in all the environments compared with the other groups (Figure 8). Cluster 4 phenotype was characterized by roots with small diameter but with many nodal roots and shallow growth angles. The second best was the root phenotype representing cluster 2, which also produced small diameter roots, but with reduced number of nodal roots and steeper growth angles. Interestingly, the phenotype of cluster 4 showed a dimorphic system with significantly greater root length in the shallow and deeper soil layers (Figure 9), surpassing the root depth of the steep angle phenotype of cluster 2. This can be explained by the shallow angles which promote a distribution of the root length in the topsoil, and by reduced root diameter, which reduces the metabolic cost of roots and promotes a deeper distribution of root length. This can be visualized when we compare the total carbon cost and root length of clusters 2 and 4. The total root carbon cost of cluster 4 is less compared with cluster 2, which produces an increased root length (Figure 10). We dissected the root phenotypes of cluster 2 and 4 to understand what were the variables that contributed to their improved performance. For cluster 2, reduced cortical cell file number alone improved shoot growth by 94, 41 and 111% under isolated drought, nitrogen and phosphorus stress (Figure 11). For cluster 4, reduced cortical cell file number improved shoot growth by 137%, 66% and 216% under isolated drought, nitrogen and phosphorus stress (Figure 11). On the other hand, we observed that small cortical cell size reduced biomass compared to the complete phenotype, indicating that the main effect of parsimonious anatomical phenotypes is coming from reduced cortical cell file number instead of cortical cell size. The contribution of other elements such as angle, nodal root number and seminal root number was irrelevant in cluster 4 (Figure 11). However, in cluster 2, reduced nodal root number contributed 8%, 17%, and 22% as increment in biomass compared to the reference (cluster 3), under drought, nitrogen and phosphorus stress, respectively. Cluster 1, which was characterized by having thick roots with steeper angles, had higher root carbon cost (Figure 10), reduced root depth (Figure 9) and reduced shoot biomass (Figure 8). Cluster 3, the average phenotype, occupied the third place in production of biomass among the 4 groups. In general, cluster 2 and 4, which had a reduced root diameter and reduced cost of soil exploration were the best performers in all environments regardless of the architecture, suggesting that phenotypes that reduce the metabolic cost are essential for broad environmental adaptation.

**Figure 8:**
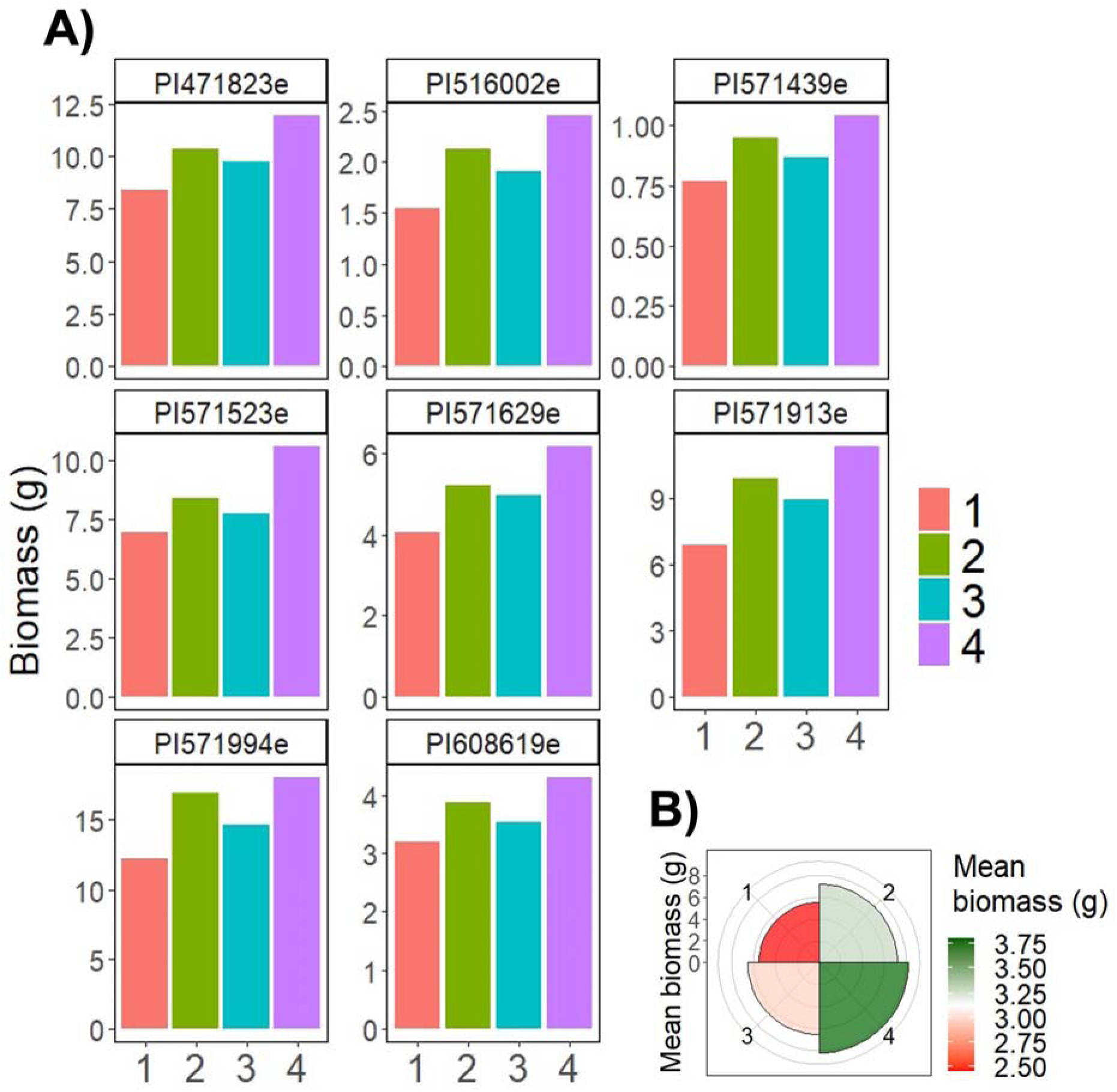
Simulation of root phenotypes of clusters 1-4 under 8 different environments of the Americas. A) Shows the simulated shoot biomass of the four clusters in every environment. The environment name corresponds location of origin from the 8 accessions previously evaluated. B) shows the mean shoot biomass of the eight environments for the root phenotypes corresponding to clusters 1, 2, 3 and 4.

**Figure 9:**
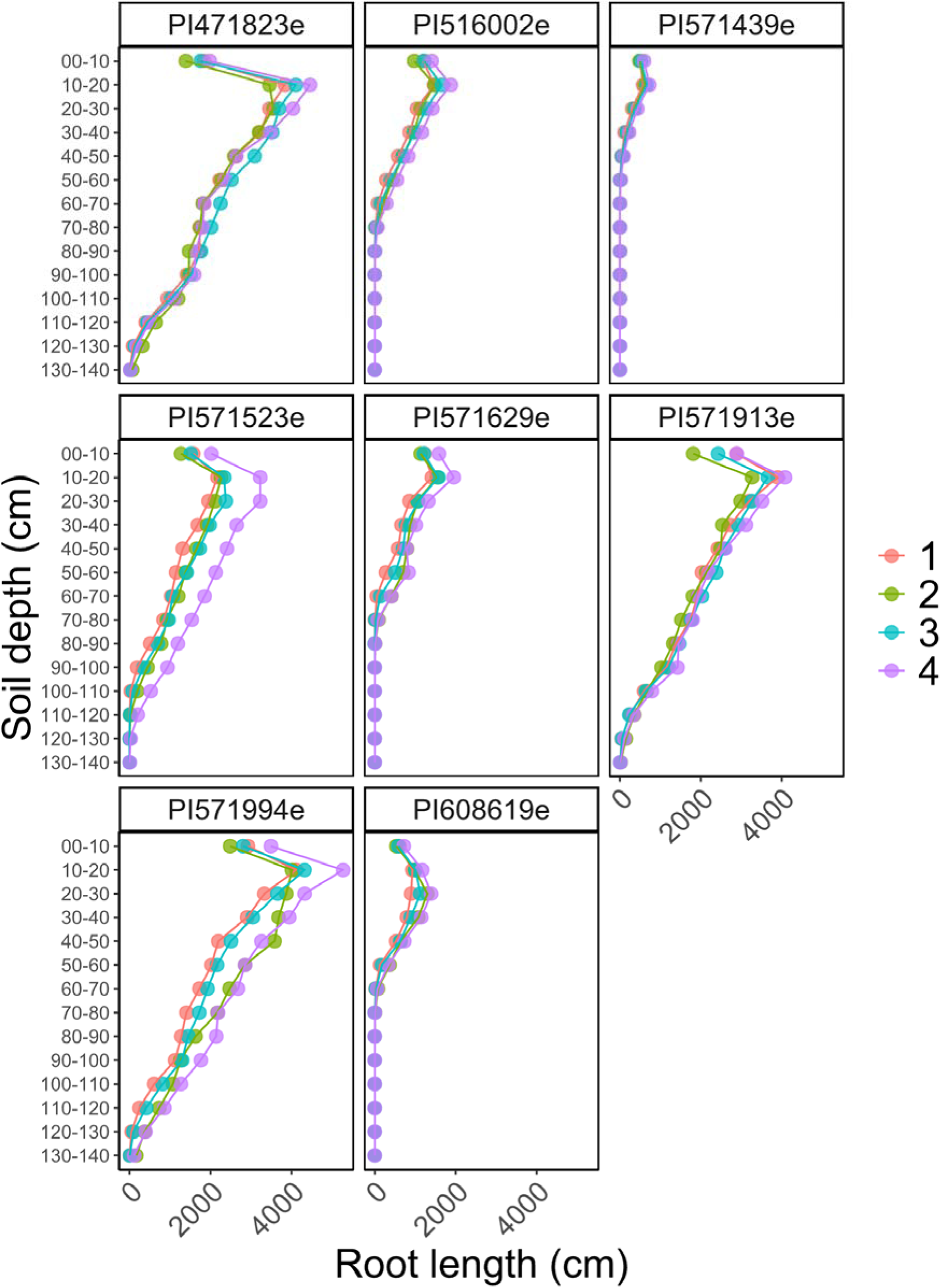
Simulated root depth of clusters 1-4 under 8 different environments. The root depth of simulations at day 40 are shown. The panel name corresponds to the environment.

**Figure 10:**
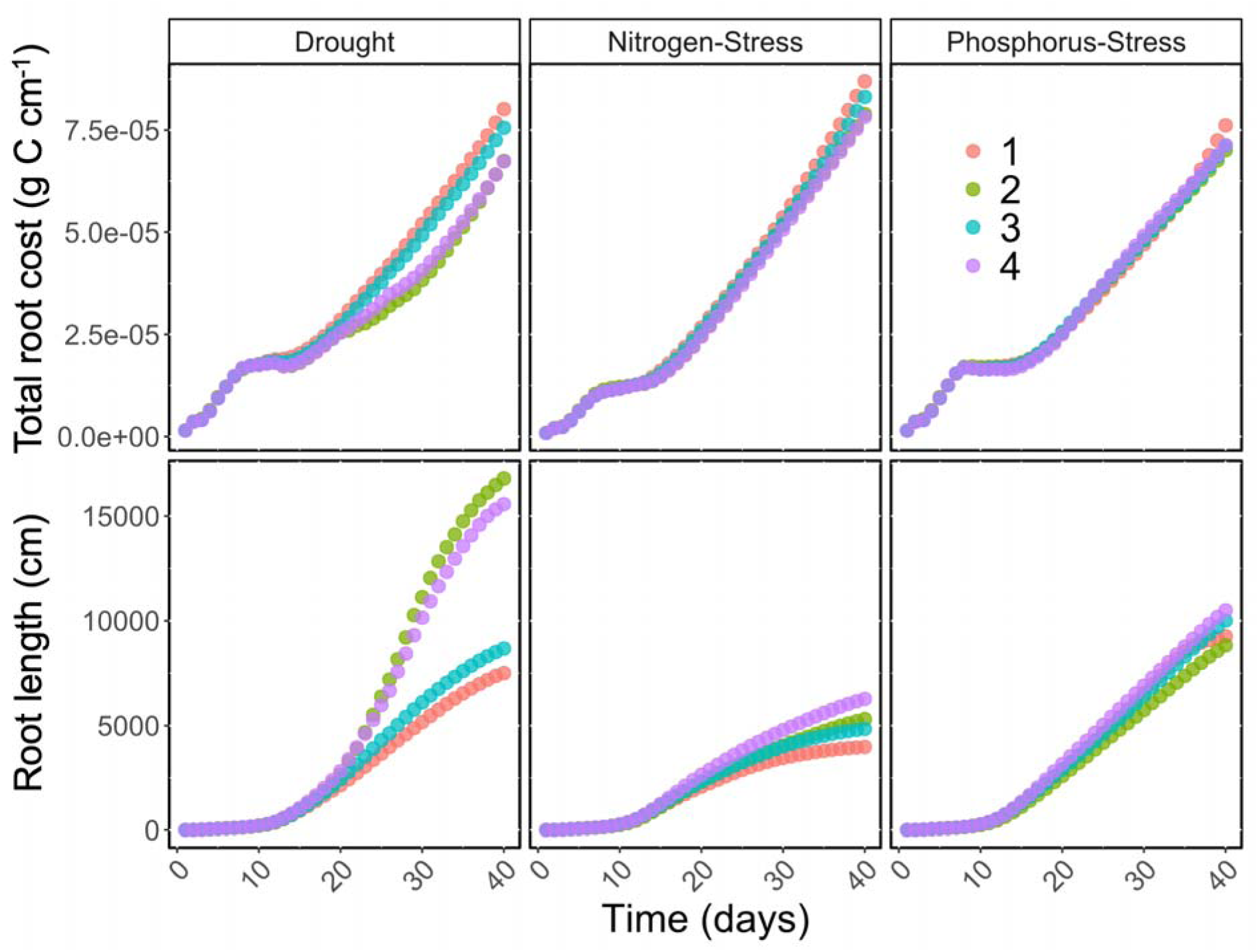
Comparison of total root carbon cost per centimeter and root length over time of phenotypic clusters 1-4 under drought, nitrogen and phosphorus stress. Different colors indicate the cluster number.

**Figure 11:**
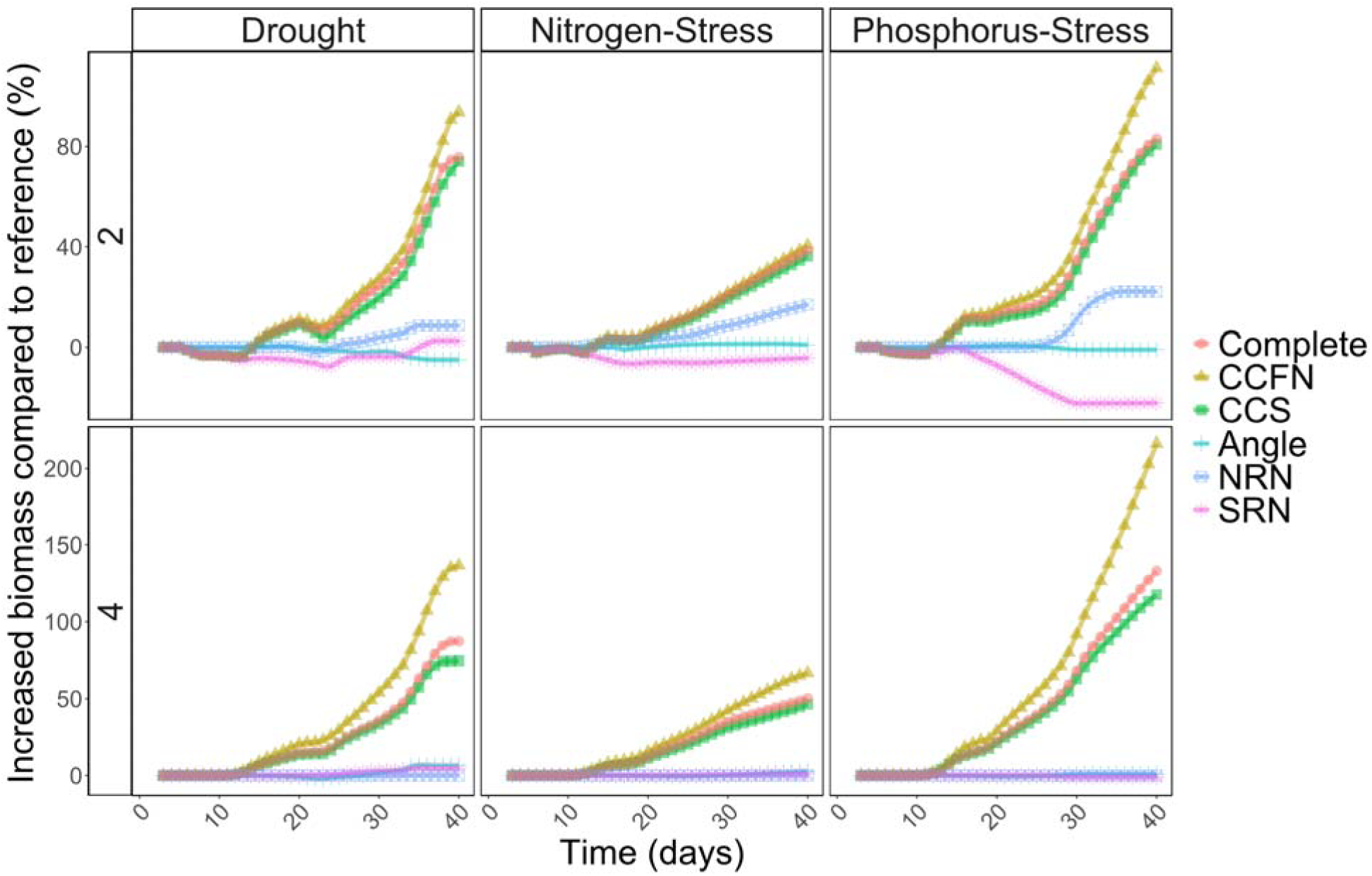
Phenotypic dissection of the root phenotypes of clusters 2 and 4 (top and bottom panel). The phenotypes of clusters 2 and 4 were deconstructed into nodal root growth angle (Angle), cortical cell file number (CCFN), cortical cell size (CCS), nodal root number (NRN), and seminal root number (SRN). Simulations correspond to 40 days. The phenotype of cluster 3 was used as a reference phenotype.

## Discussion

We explored root phenotypic variation of maize landraces from the Americas and simulated their performance across multiple environments to understand how integrated root phenotypes contribute to stability across environments. We found that non-stable accessions with steep roots and fewer nodal root numbers produced greater biomass in environments where nitrogen is the main stress factor, while accessions with shallower roots and increased number of nodal roots had better performance in environments where phosphorus was the limiting factor. Boruta feature selection showed that the most important phenotypes associated with greater biomass production across all environments are small diameters in nodes 5 and 6, large cortical cell size and reduced lateral branching frequency in node 4. We evaluated the independent effects of these phene states and found that large cortical cell size improved growth by 28%, 23 % and 114%, while reduced cortical cell file number alone improved shoot growth by 137%, 66% and 216%, under drought, nitrogen and phosphorus stress, respectively. We extended our analysis to 96 maize landraces, previously phenotyped by Burton et al., (2013), by simulating their growth with *OSR_v2* and showed that having small anatomical phenotypes improved plant growth across all environments regardless of architectural traits such as the root growth angle and number of nodal roots.

### Cheaper phenotypes offset the cost of expensive, but useful, phenotypes

Independent studies have shown that shallower root angles (Lynch 2022; Zhu et al., 2005; Miguel et al., 2015) and an increased number of nodal roots (Sun et al., 2018) colocalize roots in the phosphorus-rich topsoil, improving plant performance under phosphorus stress. In contrast, steeper root growth angles (Schneider et al., 2022; Trachsel et al., 2013) and reduced number of nodal roots (Saengwilai et al., 2014a) colocalize roots in nitrogen-rich subsoil domains and improve plant performance under nitrogen stress. Our results showed that non-stable accessions developed extreme phenotypes, for instance, shallow angles and a large number of nodal roots for PI471823, and steeper root angles with fewer nodal root numbers for PI571994. Our simulations showed that PI471823 is better adapted to environments with phosphorus stress, while PI571994 is better adapted to environments with nitrogen stress (Figure 3). Both accessions come from humid regions with relatively mild nitrogen stress (Costa Rica [PI471823], Madre de Dios, Peru [PI571994]). However, Costa Rica soils have 0.289 ppm available phosphorus while Madre de Dios soils have 0.649 ppm phosphorus (Jaramillo-Velasteguí, 2011). Since Costa Rican soils have low phosphorus bioavailability, it is logical that PI471823 presents root phenotypes optimized for phosphorus stress. It is interesting that the synthetic phenotypes with broad adaptation were characterized by having anatomical phenotypes that reduce the metabolic cost of soil exploration, while synthetic phenotypes for specific environments were characterized by anatomical phenotypes but also root architectural phenotypes such as nodal root number and root angle. This indicates that anatomical phenotypes that reduce the metabolic cost improve plant performance regardless of the environment, while root architecture phenotypes provide a more environmental specific adaptation. Our results are consistent with previous theoretical studies (Rangarajan et al., 2018; Dathe et al., 2016) and with empirical results observed in bean genotypes by Strock et al., (2019) where integrated phenotypes were associated with adaptation to low fertility and drought.

The structure-function analysis of 96 landraces showed that clusters 1 and 2 were preferentially distributed in North and South America, respectively, while clusters 3 and 4 were distributed broadly between the Americas. This reinforces the hypothesis that there are stable and non-stable root phenotypes, where the non-stable phenotypes are restricted to specific environments, while the stable phenotypes are more broadly adapted to a range of environments. The simulations of clusters 1-4 in multiple environments showed that cluster 2 (specifically from South America) and cluster 4 (broadly distributed in the Americas) were the best performers under all environments even though they showed completely contrasting root architectures. It is interesting that the clusters of stable root (broadly adapted) phenotypes were both broadly distributed in the Americas in one case (cluster 4) and coming from a particular region in another case (cluster 2). On the other hand, the non-stable (environment-specific) root phenotypes were also both broadly distributed in the Americas in one case (cluster 3), and region-specific in another case (cluster 1). This suggests that the stability of a root phenotype is not restricted to their environmental origin, indicating that root phenotypes that provide broad adaptation, such as anatomical phenotypes that reduce the metabolic cost of soil exploration, might appear equally by inheritance from previous populations locally adapted to their conditions, but also as a result of convergent evolution. However, maize landraces migrate constantly to different regions of the Americas due to seed dispersal by farmers and traders, producing a potential mismatch between the genetic relationships and the geographic distribution. Phylogenetic studies that integrate root phenotypes will be required to understand this phenomenon in more detail. Overall, our results showed that the broad adaptation of root phenotypes is not restricted to the environmental origin and might have also been the result of convergent evolution.

It has been proposed that yield stability is associated with tolerance to stress (Tollenaar and Lee, 2002). Several long-term studies have shown that yield stability increased in fields that had optimal fertilization (Ahrends et al., 2020; Knapp et al., 2018). Our results showed that the most important phene states conferring adaptation to a wide range of environments are large cortical cell size and reduced diameter of roots in nodes 5 and 6. Several studies have shown that large cortical cell size reduce the metabolic cost of soil exploration and improve plant adaptation to nitrogen stress (Lopez-Valdivia et al., 2023), phosphorus stress (Sidhu and Lynch., 2024) and drought (Chimungu et al., 2014a). Our results suggest that phenotypes that reduce the metabolic cost of soil exploration and improve plant response to stress are also associated with yield stability. It is important to note that root diameter is an aggregate phenotype composed of multiple elemental phenes such as cortical cell file number, cortical cell size, and stelar phenes. Cortical cell diameter is a better predictor of the energy cost of the root compared with the root diameter as it considers the living cortical area (Colombi et al., 2019; Jaramillo et al., 2013). Additionally, a functional-structural analysis of 96 landraces showed 4 main clusters of root phenotypes. Our results showed that clusters 2 and 4, with a small root cross section area (RXSA), reduced stele area (TSA), reduced cortical cell file number X.CF, and reduced cortical cell area (CCA), showed a reduced total carbon cost (Figure 10). This reduction in total root carbon cost was associated with increased total root length (Figure 10), which is translated to improved soil exploration and better performance across the eight simulated environments even though they had contrasting phenotypes for root growth angle and number of nodal roots. Among all phenes of the 96 landraces, reduced cortical cell file number had the greatest contribution to simulated biomass production under isolated stress for drought, nitrogen and phosphorus. The improved performance of clusters 2 and 4, with contrasting root architecture, suggests that the potential disadvantages of the extreme root architecture (e.g. very shallow or very steep), will be compensated by a reduced cost of soil exploration due to small anatomical phenotypes. Overall, our results suggest that anatomical phenes that reduce the metabolic cost of soil exploration are associated with greater yield stability.

It has been shown that increased lateral branching increases root respiration and root maintenance cost (Zhan et al 2015; Zhan and Lynch, 2015). Our feature selection prediction showed that reduced lateral branching frequency in node 4 would be important for general adaptation across environments (Table 2). However, when we tested the independent effects of each phene state, lateral branching frequency did not confer a significant benefit under nitrogen and phosphorus stress. This can be explained by the relative utility of this phene. For instance, reduced branching frequency was also selected by Boruta for specific synthetic phenotypes for environments PI608619e, PI571439e, PI571629e, PI571994e (Table 1). It is important to note that we are testing the effect of lateral branching frequency in node 4 only, while previous studies showed a strong effect on lateral branching frequency when they were evaluated across all axial roots (Postma et al., 2014; Rangarajan et al., 2018). Further analysis will be needed to determine the effect of lateral branching frequency in each of these environments. This is interesting because it might reflect the tradeoffs between the ability of a phenotype to produce greater biomass in a wide variety of environments compared with the ability to produce biomass in its optimal environment (Weiner et al., 2021).

Our results support the hypotheses that root anatomical phenotypes that reduce the metabolic cost of soil exploration and intermediate root architectures will be useful targets of selection for crops with greater yield stability. Previous studies explored the fitness landscape utilizing massive arrays of possible root phenotypes (Ajmera et al., 2022; Rangarajan et al., 2018), or more refined approaches such as multi-objective evolutionary algorithms (Rangarajan et al., 2022) to identify integrated phenotypes adapted to diverse environments. Our work is the first study that explores the utility of different integrated root systems of maize landraces from the Americas across multiple environments *in silico*. This is an example of how functional-structural plant/soil models can be used to identify optimal and stable integrated phenotypes starting from a small population of plants.

## Abbreviations

OSRv2: OpenSimRoot_v2
BF: lateral branching frequency
CCS: cortical cell size
Dia: diameter
AN: root angle
N#: node number

## Literature Cited

1. Ajmera, I., Henry, A., Radanielson, A. M., Klein, S. P., Ianevski, A., Bennett, M. J., Band, L. R., & Lynch, J. P. (2022). Integrated root phenotypes for improved rice performance under low nitrogen availability. Plant Cell and Environment 45, 805–822.

2. Alm, D. M., Cavelier, J., and Nobel, P. S. (1992). A finite-element model of radial and axial conductivities for individual roots: Development and validation for two desert succulents. Annal of Botany 69, 87–87.

3. Burridge, J. D., Findeis, J. L., Jochua, C. N., Miguel, M. A., Mubichi-Kut, F. M., Quinhentos, M. L., Xerinda, S. A., & Lynch, J. P. (2019). A case study on the efficacy of root phenotypic selection for edaphic stress tolerance in low-input agriculture: Common bean breeding in Mozambique. Field Crops Research 244, 1 December 2019, 107612.

4. Burton, A., Brown, K. M., & Lynch, J. P. (2013). Phenotypic diversity of root anatomical and architectural traits in Zea species. Crop Science 53, 1042–1055.

5. Chimungu, J. G., Brown, K. M., & Lynch, J. P. (2014a). Large root cortical cell size improves drought tolerance in maize. Plant Physiology 166, 2166–2178.

6. Chimungu, J. G., Brown, K. M., & Lynch, J. P. (2014b). Reduced root cortical cell file number improves drought tolerance in maize. Plant Physiology 166, 1943–1955.

7. Chimungu, J. G., Loades, K. W., & Lynch, J. P. (2015b). Root anatomical phenes predict root penetration ability and biomechanical properties in maize (*Zea Mays*). Journal of Experimental Botany 66, 3151–3162.

8. Chimungu, J. G., Maliro, M. F., Nalivata, P., Kanyama-Phiri, G., Brown, K., & Lynch, J. (2015a). Utility of root cortical aerenchyma under water limited conditions in tropical maize (*Zea mays L*.). Field Crops Research 171, 86–98.

9. Colombi, T., Herrmann, A. M., Vallenback, P., & Keller, T. (2019). Cortical cell diameter is key to energy costs of root growth in wheat. Plant Physiology 180, 2049–2060.

10. Dathe, A., Postma, J. A., Postma-Blaauw, M. B., & Lynch, J. P. (2016). Impact of axial root growth angles on nitrogen acquisition in maize depends on environmental conditions. Annals of Botany 118, 401–414.

11. Doussan, C., Pagès, L., and Vercambre, G. (1998). Modelling of the hydraulic architecture of root systems: an integrated approach to water absorption–model description. Annals of Botany 81, 213–223.

12. Fan, X. M., Kang, M. S., Chen, H., Zhang, Y., Tan, J., & Xu, C. (2007). Yield stability of maize hybrids evaluated in multi-environment trials in Yunnan, China. Agronomy Journal 99, 220–228.

13. Francis, T. R., & Kannenberg, L. W. (1978). Yield stability studies in short-season maize: Relationship to plant-to-plant variability. Canadian Journal of Plant Science 58, 1035–1039.

14. Galindo-Castañeda, T., Brown, K. M., & Lynch, J. P. (2018). Reduced root cortical burden improves growth and grain yield under low phosphorus availability in maize. Plant Cell and Environment 41, 1579–1592.

15. Gao, Y., & Lynch, J. P. (2016). Reduced crown root number improves water acquisition under water deficit stress in maize (Zea mays L.). Journal of Experimental Botany 67, 4545–4557.

16. Hochholdinger, F., Yu, P., & Marcon, C. (2018). Genetic control of root system development in maize. Trends in Plant Science 23, 79–88.

17. Jaramillo-Velasteguí, R. E. (2011). The edaphic control of plant response to climate change: Extent, interactions, and mechanisms of plant adaptation. [Doctoral dissertation, The Pennsylvania State University].

18. Jaramillo, R. E., Nord, E. A., Chimungu, J. G., Brown, K. M., & Lynch, J. P. (2013). Root cortical burden influences drought tolerance in maize. Annals of Botany 112, 429–437.

19. Jia, X., Liu, P., & Lynch, J. P. (2018). Greater lateral root branching density in maize improves phosphorus acquisition from low phosphorus soil. Journal of Experimental Botany, 69, 4961–4790.

20. Jia, X., Wu, G., Strock, C., Li, L., Dong, S., Zhang, J., Zhao, B., Lynch, J. P., & Liu, P. (2022). Root anatomical phenotypes related to growth under low nitrogen availability in maize (*Zea mays L*.). Plant and Soil 474, 265–276.

21. Lopez-Valdivia, I., Yang, X., & Lynch, J. P. (2023). Large root cortical cells and reduced cortical cell files improve growth under suboptimal nitrogen in silico. Plant Physiology 192, 2261–2275.

22. Lynch, J. P. (2013). Steep, cheap, and deep: An ideotype to optimize water and N acquisition by maize root systems. Annals of Botany 112, 347–357.

23. Lynch, J. P. (2022). Harnessing root architecture to address global challenges. Plant Journal 109, 415–431.

24. Lynch, J. P., & Brown, K. M. (2001). Topsoil foraging – an architectural adaptation of plants to low phosphorus availability. Plant and Soil 237, 225–237.

25. Lynch, J. P., Galindo-Castañeda, T., Schneider, H. M., Sidhu, J. S., Rangarajan, H., & York, L. M. (2023). Root phenotypes for improved nitrogen capture. Plant and Soil 10.1007/s11104-023-06301-2.

26. Lynch, J. P., Mooney, S. J., Strock, C. F., & Schneider, H. M. (2022). Future roots for future soils. Plant Cell and Environment 45, 620–636.

27. Lynch, J. P., Strock, C. F., Schneider, H. M., Sidhu, J. S., Ajmera, I., Galindo-Castañeda, T., Klein, S. P., & Hanlon, M. T. (2021). Root anatomy and soil resource capture. Plant and Soil 466, 21–63.

28. Miguel, M. A., Postma, J. A., & Lynch, J. P. (2015). Phene synergism between root hair length and basal root growth angle for phosphorus acquisition. Plant Physiology 167, 1430–1439.

29. Monteith, J. L. (1965). “Evaporation and environment,” in Symposia of the Society for Experimental Biology, vol. 19. (Cambridge University Press, Cambridge), 205–234.

30. Penman, H. L. (1948). Natural evaporation from open water, bare soil and grass. Proc. R. Soc. London. Ser. A. Math. Phys. Sci. 193, 120–145.

31. Poggio, L., de Sousa, L. M., Batjes, N. H., Heuvelink, G. B. M., Kempen, B., Ribeiro, E., & Rossiter, D. (2021). SoilGrids 2.0: Producing soil information for the globe with quantified spatial uncertainty. SOIL 7, 217–240.

25. Postma, J. A., & Lynch, J. P. (2012). Complementarity in root architecture for nutrient uptake in ancient maize/bean and maize/bean/squash polycultures. Annals of Botany, 110, 521–534.

26. Postma, J. A., Dathe, A., & Lynch, J. P. (2014). The optimal lateral root branching density for maize depends on nitrogen and phosphorus availability. Plant Physiology 166, 590–602.

27. Postma, J. A., Kuppe, C., Owen, M. R., Mellor, N., Griffiths, M., Bennett, M. J., Lynch, J. P., & Watt, M. (2017). OpenSimRoot: widening the scope and application of root architectural models. New Phytologist 215, 1274–1286.

28. Postma, J. A., Kuppe, C., Owen, M. R., Mellor, N., Griffiths, M., Bennett, M. J., Lynch, J. P., & Watt, M. (2017). OpenSimRoot: widening the scope and application of root architectural models. New Phytologist 215, 1274–1286.

29. R Core Team. (2019). R: A language and environment for statistical computing. R Foundation for Statistical Computing.

30. Rangarajan, H., Hadka, D., Reed, P., & Lynch, J. P. (2022). Multi-objective optimization of root phenotypes for nutrient capture using evolutionary algorithms. Plant Journal 111, 38–53.

31. Rangarajan, H., Postma, J. A., & Lynch, J. P. (2018). Co-optimization of axial root phenotypes for nitrogen and phosphorus acquisition in common bean. Annals of Botany 122, 485–499.

32. Rasmusson, D. C. (1987). An evaluation of ideotype breeding. Crop Science, 27, 1140–1146.

33. Saengwilai, P., Nord, E. A., Chimungu, J. G., Brown, K. M., & Lynch, J. P. (2014b). Root cortical aerenchyma enhances nitrogen acquisition from low-nitrogen soils in maize. Plant Physiology 166, 726–735.

34. Saengwilai, P., Tian, X., & Lynch, J. P. (2014a). Low crown root number enhances nitrogen acquisition from low-nitrogen soils in maize. Plant Physiology 166, 581–589.

35. Schäfer, E. D., Ajmera, I., Farcot, E., Owen, M. R., Band, L. R., & Lynch, J. P. (2022). In silico evidence for the utility of parsimonious root phenotypes for improved vegetative growth and carbon sequestration under drought. Frontiers in Plant Science 13, 1–30.

36. Schneider, H. M., Lor, V. S. N., Hanlon, M. T., Perkins, A., Kaeppler, S. M., Borkar, A. N., Bhosale, R., Zhang, X., Rodriguez, J., Bucksch, A., Bennett, M. J., Brown, K. M., & Lynch, J. P. (2022). Root angle in maize influences nitrogen capture and is regulated by calcineurin B-like protein (CBL)-interacting serine/threonine-protein kinase 15 (ZmCIPK15). Plant Cell and Environment 45, 837–853.

37. Schneider, H. M., Postma, J. A., Wojciechowski, T., Kuppe, C., & Lynch, J. P. (2017). Root cortical senescence improves growth under suboptimal availability of N, P, and K. Plant Physiology 174, 2333–2347.

38. Schneider, H. M., Strock, C. F., Hanlon, M. T., Vanhees, D. J., Perkins, A. C., Ajmera, I. B., Sidhu, J. S., Mooney, S. J., Brown, K. M., & Lynch, J. P. (2021). Multiseriate cortical sclerenchyma enhance root penetration in compacted soils. Proceedings of the National Academy of Sciences, 118(e2012087118).

39. Schneider, H. M., Yang, J. T., Brown, K. M., & Lynch, J. P. (2021). Nodal root diameter and node number in maize (*Zea mays L*.) interact to influence plant growth under nitrogen stress. Plant Direct 5, 10.1002/pld3.310.

40. Shiri, M. R. (2013). Grain yield stability analysis of maize (*Zea mays L*.) hybrids under different drought stress conditions using GGE biplot analysis. Crop Breeding Journal 3, 107–112.

41. Sidhu, J. S., & Lynch, J. P. (2024). Cortical cell size regulates root metabolic cost. The Plant Journal.118, 1343–1357

49. Simunek, J., Huang, K., & Genuchten, M. (1995). The SWMS 3D code for simulating water flow and solute transport in three-dimensional variably-saturated media. U.S. Salinity Laboratory Agricultural Research Service.

43. Sosa, J. M., Huber, D. E., Welk, B., & Fraser, H. L. (2014). Development and application of 32 MIPARTM: A novel software package for two- and three-dimensional microstructural characterization. Integr. Mater. Manuf. Innov., 3, 10.

44. Strock, C. F., Burridge, J., Massas, A. S. F., Beaver, J., Beebe, S., Camilo, S. A., Fourie, D., Jochua, C., Miguel, M., Miklas, P. N., Mndolwa, E., Nchimbi-Msolla, S., Polania, J., Porch, T. G., Rosas, J. C., Trapp, J. J., & Lynch, J. P. (2019). Seedling root architecture and its relationship with seed yield across diverse environments in *Phaseolus vulgaris*. Field Crops Research 237, 53–64.

45. Strock, C. F., Rangarajan, H., Black, C. K., Schäfer, E. D., & Lynch, J. P. (2022). Theoretical evidence that root penetration ability interacts with soil compaction regimes to affect nitrate capture. Annals of Botany 129, 315–330.

53. Strock, C. F., Schneider, H. M., Galindo-Castañeda, T., Hall, B. T., van Gansbeke, B., Mather, D. E., Roth, M. G., Chilvers, M. I., Guo, X., Brown, K., & Lynch, J. P. (2019). Laser ablation tomography for visualization of root colonization by edaphic organisms. Journal of Experimental Botany 70, 5327–5342.

47. Sun, B., Gao, Y., & Lynch, J. P. (2018). Large crown root number improves topsoil foraging and phosphorus acquisition. Plant Physiology 177, 90–104.

48. Tollenaar, M., & Lee, E. A. (2002). Yield potential, yield stability, and stress tolerance in maize. Field Crops Research 75, 161–169.

49. Trachsel, S., Kaeppler, S. M., Brown, K. M., & Lynch, J. P. (2011). Shovelomics: High throughput phenotyping of maize (*Zea mays L.*) root architecture in the field. Plant and Soil 341, 75–87.

50. Trachsel, S., Kaeppler, S. M., Brown, K. M., & Lynch, J. P. (2013). Maize root growth angles become steeper under low N conditions. Field Crops Research 140, 18–31.

51. USDA, Agricultural Research Service, National Plant Germplasm System. (2024). Germplasm Resources Information Network (GRIN Taxonomy). National Germplasm Resources Laboratory, Beltsville, Maryland.

52. Venables, W.N., Ripley, B.D. (2002). Modern Applied Statistics with S, Fourth edition. (Springer, New York).

53. Weiner, J., Du, Y. L., Zhao, Y. M., & Li, F. M. (2021). Allometry and yield stability of cereals. Frontiers in Plant Science, 12, 1–5. doi: 10.3389/fpls.2021.681490

54. Williams, A., Hunter, M. C., Kammerer, M., Kane, D. A., Jordan, N. R., Mortensen, D. A., Smith, R. G., Snapp, S., & Davis, A. S. (2016). Soil water holding capacity mitigates downside risk and volatility in US rainfed maize: Time to invest in soil organic matter? PloS ONE 11, 1–11.

55. Wickham, H. ggplot2: Elegant Graphics for Data Analysis (Springer, New York, 2016).

56. Yang HS, Janssen BH. (2000). A mono-component model of carbon mineralization with a dynamic rate constant. European Journal of Soil Science 51, 517–529.

57. Zhan, A., & Lynch, J. P. (2015). Reduced frequency of lateral root branching improves N capture from low-N soils in maize. Journal of Experimental Botany 7, 2055–2065.

58. Zhan, A., Schneider, H., & Lynch, J. P. (2015). Reduced lateral root branching density improves drought tolerance in maize. Plant Physiology 168, 1603–1615.

47. Zhao, J., & Yang, X. (2018). Distribution of high-yield and high-yield-stability zones for maize yield potential in the main growing regions in China. Agricultural and Forest Meteorology 248, 511–517.

48. Zhu, J., Brown, K. M., & Lynch, J. P. (2010b). Root cortical aerenchyma improves the drought tolerance of maize (*Zea mays L*.). Plant, Cell and Environment 33, 740–749.

49. Zhu, J., Kaeppler, S. M., & Lynch, J. P. (2005). Topsoil foraging and phosphorus acquisition efficiency in maize (*Zea mays*). Functional Plant Biology 32, 749–762.

50. Zhu, J., Zhang, C., & Lynch, J. P. (2010a). The utility of phenotypic plasticity of root hair length for phosphorus acquisition. Functional Plant Biology 37, 313–322.

