## Supplementary data for "Exploring yield stability and the fitness landscape of maize landrace root phenotypes *in silico*"

**This pdf includes:**

Table S1

Table S1: Origin information of the eight accessions used in this study.

| Accession | Name | Origin | Coordinates | Climatic Zone |
| --- | --- | --- | --- | --- |
| PI 471823 | Costa Rica 139 | Costa Rica | 10.88518100, -85.00984500 | Humid |
| PI 516002 | ARZM 16 001 | Mendoza, Argentina | -33.26666667, -68.15000000 | Arid |
| PI 571439 | Lambayeque 151 | Lambayeque, Peru | -6.70000000, -79.91666667 | Arid |
| PI 571523 | Ica 8 | Ica, Peru | -14.18333333, -75.73333333 | Arid |
| PI 571629 | Cuzco 22 | Cusco, Peru | -12.81666667, -72.71666667 | Sub-humid |
| PI 571913 | Huanuco 191 | Huánuco, Peru | -10.06666667, -75.53333333 | Humid |
| PI 571994 | Madre de Dios 22 | Madre de Dios, Peru | -13.68333333, -69.53333333 | Humid |
| PI 608619 | Minnesota 13 | Colorado, United States | 39.16666667, -108.80000000 | Arid |
